# Adaptive Cortical Parcellations for Source Reconstructed EEG/MEG Connectomes

**DOI:** 10.1101/097774

**Authors:** Seyedeh-Rezvan Farahibozorg, Richard N Henson, Olaf Hauk

**Author notes:** Corresponding author: Seyedeh-Rezvan Farahibozorg, MRC Cognition and Brain Sciences Unit, University of Cambridge, 15 Chaucer Road, Cambridge, UK, CB2 7EF.

## Abstract

There is growing interest in the rich temporal and spectral properties of the brain's functional connectome that are provided by Electro- and Magnetoencephalography (EEG/MEG). However, the problem of leakage between brain sources that arises when reconstructing brain activity from EEG/MEG recordings outside the head makes it difficult to distinguish true connections from spurious connections, even when connections are based on measures that ignore zero-lag dependencies. In particular, standard anatomical parcellations for potential cortical sources tend to over- or under-sample the real spatial resolution of EEG/MEG. By using information from cross-talk functions (CTFs) that objectively describe leakage for a given sensor configuration and distributed source reconstruction method, we introduce methods for optimising the number of parcels while simultaneously minimising the leakage between them. More specifically, we compare two image segmentation algorithms: 1) a split-and-merge (SaM) algorithm based on standard anatomical parcellations and 2) a region growing (RG) algorithm based on all the brain vertices with no prior parcellation. Interestingly, when applied to minimum-norm reconstructions for EEG/MEG configurations from real data, both algorithms yielded approximately 70 parcels despite their different starting points, suggesting that this reflects the resolution limit of this particular sensor configuration and reconstruction method. Importantly, when compared against standard anatomical parcellations, resolution matrices of adaptive parcellations showed notably higher sensitivity and distinguishability of parcels. Furthermore, extensive simulations of realistic networks under various circumstances revealed significant improvements in network reconstruction accuracies, particularly in reducing false leakage-induced connections. Adaptive parcellations therefore allow a more accurate reconstruction of functional EEG/MEG connectomes.

**Highlights:**

- We introduce adaptive cortical parcellation algorithms for E/MEG source estimation.
- Algorithms are based on cross-talk functions and image segmentation methods.
- The resulting parcellations yielded ~70 parcels regardless of starting point.
- Sensitivity and distinguishability improved compared to anatomical parcellations.
- Accuracy of realistic whole-brain network reconstruction improved significantly.

## 1 Introduction

Connectivity analyses of source estimated Electro- and Magnetoencephalography (EEG/MEG) can provide a millisecond-by-millisecond map of functional and effective interactions (Bastos & Schoffelen 2016; Greenblatt et al. 2012) among multiple brain areas in resting state as well as during task performance (Brookes et al. 2016; Colclough et al. 2016; Palva et al. 2010). Consequently, there has been growing interest in reconstructing the human brain connectome to obtain time- and frequency-resolved whole-brain networks (Palva & Palva 2012). Studies on anatomical and functional MRI connectomics have revealed important properties of the brain in health and disease, particularly concerning changes in “hubs” and the associated “rich club” of highly-connected regions (Bullmore & Sporns 2009; Crossley et al. 2014; van den Heuvel & Sporns 2011). The growing field of EEG/MEG connectomics is anticipated to take this approach further by vastly increasing the temporal and spectral resolution of the human connectome (Brookes et al. 2011; de Pasquale et al. 2010). However, the spatial resolution of EEG/MEG data is limited, because several thousand sources of activation in the brain must be estimated from maximally a few hundred sensor recordings.

The limited spatial resolution causes the so-called leakage or cross-talk problem for linear and linearly constrained distributed EEG/MEG source estimation: activity estimated in one region of interest (ROI) can be affected by leakage from locations outside this ROI, possibly including locations at large distances (Lachaux et al. 1999; Schoffelen & Gross 2009; Hauk et al. 2011). This poses serious challenges for the interpretation of connectivity results, since increased connectivity between two ROIs may not only be caused by true connections between the time courses of these ROIs, but also by signals leaked into these ROIs from other brain locations, thus leading to spurious connectivity findings (Colclough et al. 2015). This is particularly important for the estimation of whole-brain connectivity and applications of graph theoretical measures. For example, one ROI in a network may be identified as a hub (i.e. showing strong connections to several other ROIs) if it receives strong leakage from multiple other ROIs.

Most previous EEG/MEG studies have adopted parcellations from anatomical or fMRI research for whole-brain connectivity analysis (Colclough et al. 2016; Brookes et al. 2016; Tewarie et al. 2016). Some studies have orthogonalised source-reconstructed timeseries across parcels, in order to remove any zero-lag correlation, such as that induced by leakage (Brookes et al. 2012; Hipp et al. 2012; Colclough et al. 2015). While this may be suitable if connectivity is estimated from more slowly-varying amplitude envelopes of ongoing oscillatory activity, it also potentially removes true zero-lag connectivity that is not an artefact of cross-talk. Additionally, considering the spatial resolution of EEG/MEG, anatomical parcellations may not be optimal and recent studies have suggested that EEG/MEG-based parcellations can be more informative (Brookes et al. 2016). The ideal parcellation should be sensitive to as much of the cortex as possible, with each parcel having high sensitivity to activity arising from itself, and low leakage from other parcels. CTFs can be used to characterise leakage among different brain areas (Liu et al. 1998; Hauk et al. 2011). Some previous studies have suggested using CTFs to minimise leakage between a small number of ROIs. Wakeman (2013), for example, sub-selected a number of vertices as representative for each of a few ROIs that had minimal cross-talk with the other ROIs, while Hauk and Stenroos (2014) proposed a method that optimises spatial filters for source reconstruction in order to produce zero cross-talk among a small set of brain sources and minimal cross-talk from other sources.

While these methods are optimised for the case of few spatially distinct sources, their extension to whole-brain connectivity analysis is limited. Palva et al. (2010) introduced a parcellation for graph theoretical analysis of single subject data by taking into account the source-sensor geometry of EEG/MEG. They used a clustering algorithm to parcellate the cortex into 365 patches (equal to the number of sensors), based on phase synchrony patterns estimated from simulated data generated from white noise in source space. Korhonen et al. (2014) introduced sparse weights to collapse the source space based on the forward and inverse modelling of simulated noise in the source space. Their method aims at assigning optimal vertices to a fixed set of parcels and extracting the parcel time course as a weighted sum of the assigned vertices. This method utilises phase coherence between the true and estimated sources in order to maximise the fidelity of assigned vertices to the recipient parcel. Unlike the aforementioned Palva et al. method (2010), the sparse weights approach is suitable for group as well as single subject analysis and is based on the anatomical parcellations. However, while the sparse weights approach provides a way of extracting parcel time courses based on the spatial limitations of EEG/MEG, obtaining an adaptive parcellation that can optimise both the number and location of parcels, as well as vertex selection within those parcels, has remained a challenge (Korhonen et al. 2014; Bullmore & Bassett 2011). This is in spite of the fact that considering the fast growing field of connectomics, obtaining accurate parcellations of the brain (Glasser et al. 2016; Wang et al. 2015) has become desirable and consequently accurate and adaptive parcellations of the cortex for EEG/MEG data should prove useful.

Here, we utilise CTFs as a direct measure of spatial leakage to address the limitations of the aforementioned methods systematically. For this purpose, we have implemented two CTF-informed image segmentation algorithms (Gonzalez & Woods 2007) that parcellate the cortical surface into the maximum number of distinguishable parcels. In the first approach, we started from standard anatomical parcellations and modified the parcels using a CTF-informed split-and-merge (SaM) algorithm. The main idea is to merge parcels that produce highly overlapping CTFs, split parcels that produce distinguishable patterns of cross-talk, remove parcels for which EEG/MEG show low sensitivity, and identify, for each parcel, a group of representative vertices that show high sensitivity and specificity to that particular parcel relative to the rest of the brain. This approach is suitable for studies that require a particular anatomical labelling of parcels. In the second approach, we start from all the brain vertices with no prior parcellation. A CTF-informed region growing algorithm is used to create parcels around the vertices that show highest sensitivity and specificity of CTFs on the cortex. These parcels are then optimised with respect to specificity and sensitivity using a SaM algorithm. This approach should prove useful for studies where no strict anatomical labels are required.

Both algorithms yield adaptive parcellations since CTF patterns may change depending on the choice of head models, inverse operators, measurement configurations (i.e. EEG, MEG or their combination) and signal-to-noise ratios (SNR) of the data. Additionally, the proposed algorithms can use data from multiple subjects and yield parcellations suitable for group analysis through morphing the cortical surfaces from single subjects to a standard average space (e.g. MNI space). We evaluate the performance of the proposed algorithms by measuring the sensitivity and specificity of the CTFs of the final parcels to themselves as compared to the rest of the brain, and comparing performance to those of two standard anatomical atlases in the Freesurfer software (Desikan-Killiany (Desikan et al. 2006) and Destrieux (Destrieux et al. 2010)). Furthermore, we compared the performance of different parcellations by means of spectral connectivity analysis of simulated event-related networks in source space, and under various conditions in terms of number and locations of active sources, percentage of connections among the sources and realistic SNRs of the data. We show that EEG/MEG-adaptive parcellations result in a more accurate network reconstruction for both zero-lag and non-zero-lag connectivity metrics.

## 2 Theory

### 2.1 EEG/MEG source estimation and spatial resolution

In this section we introduce the concepts of the resolution matrix and cross-talk functions, which are the basis for the parcellation algorithms described in later Methods section.

#### 2.1.1 EEG/MEG forward and inverse solution

In forward modelling of EEG/MEG data, assuming a linear relationship between data and sources, the leadfield matrix (**G**) maps the sources of activity on the cortex to the electric and magnetic signals measured using EEG and MEG sensors (Hämäläinen & Ilmoniemi. 1994). Therefore, signal at each sensor is modelled as a weighted sum of the activities of all the sources in the brain:

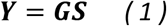

where **Y** is an N_ch_ ⨯ N_t_ matrix of the measured signal at the sensor locations, the time-invariant matrix **G** denotes the leadfield of size N_ch_⨯N_s_ and **S** denotes the source activity matrix which is of size N_s_ ⨯ N_t_ (Nch: number of recording channels, N_t_: number of time points, N_s_: number of sources).

For EEG/MEG, linear source estimation methods are often employed in order to obtain a solution for **S** in Equation 1, i.e. if **D= *Y* + ϵ** is the matrix of the measured data of size N_ch_⨯N_t_ (which contains activity from brain sources in Equation 1 plus noise), the source activity is estimated as:

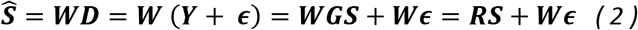

where **W** is the inverse operator of size N_s_⨯N_ch_ that maps measurements to the sources, **Ŝ** is the matrix of estimated sources of size N_s_ ⨯ N_t_ and **ԑ** denotes the measurement noise matrix of size N_ch_ ⨯ N_t_ and **R=WG** is the resolution matrix.

#### 2.1.2 Resolution matrix and CTFs

In Equation 2, the resolution matrix **R** can be used to quantify the relationship between true and estimated sources. The diagonal elements of **R** indicate the sensitivity of each estimated source to itself, and off-diagonal elements quantify the degree to which estimated sources are affected by the signal from all other sources in the brain (Grave De Peralta Menendez et al. 1997; Liu et al. 1998). An accurate estimation of source activity in the brain would for example be possible if **G** was a full-ranked square matrix (i.e. equal number of sensors and sources) and in the absence of measurement noise. In such an ideal scenario **W** would be the inverse of **G**, **R = G**^**-1**^**G = I** would be an identity matrix and the estimated sources would precisely match the true sources. However, the EEG/MEG inverse problem is a highly underdetermined problem and the resolution matrix has non-zero off-diagonal elements. These off-diagonal elements introduce the leakage or cross-talk in the EEG/MEG inverse solutions. One method of estimating the inverse operator is L2 minimum norm estimates (MNE):

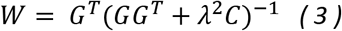

Where λ is the regularisation parameter and C is the noise covariance of the data. According to Backus and Gilbert (Backus & Gilbert 1970), λ provides a trade-off between spatial resolution and stability for the source estimate. Consequently, the resolution matrix for the L2 MNE will be obtained as:

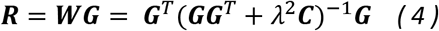

It is worth noting that the i^th^ row of **R** describes the cross-talk from all sources in the brain into the estimate for activity of the i^th^ source. These rows have therefore been called cross-talk functions (CTFs) (Liu et al. 1998; Hauk et al. 2011). Therefore, the cross-talk from the j^th^ to the i^th^ source is defined as:

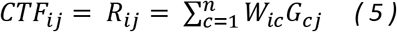

where n is the number of sensors in the brain. As explained above, ideally *R_ij_* should be 0 for any i≠j and 1 for i=j. If an element *R_ij_* is zero, there is no cross-talk from the j^th^ source into the estimate for the i^th^ source. If two CTFs are largely non-overlapping, this means they are sensitive to different areas of the brain. If *R_ij_* is much larger than the value of *R_ik_* (k being a third source in the brain), this means that the estimator is more prone to receive cross-talk from the j^th^ source than from the k^th^ source. Note that a CTF is necessarily a linear combination of the leadfields (i.e. rows of **G**). Therefore, CTFs cannot be designed to take on any arbitrary shape, but are constrained by the measurement configuration. Therefore, CTFs offer a direct way of quantifying the cross-talk problem for linear estimators of a given measurement configuration, which can be used to find an optimal parcellation of the source space based on objective criteria.

#### 2.1.3 Using CTFs to modify anatomical atlases

Two main problems can arise from utilising anatomical parcellations with EEG/MEG, which we illustrate in Fig. 1:

**Figure 1:**
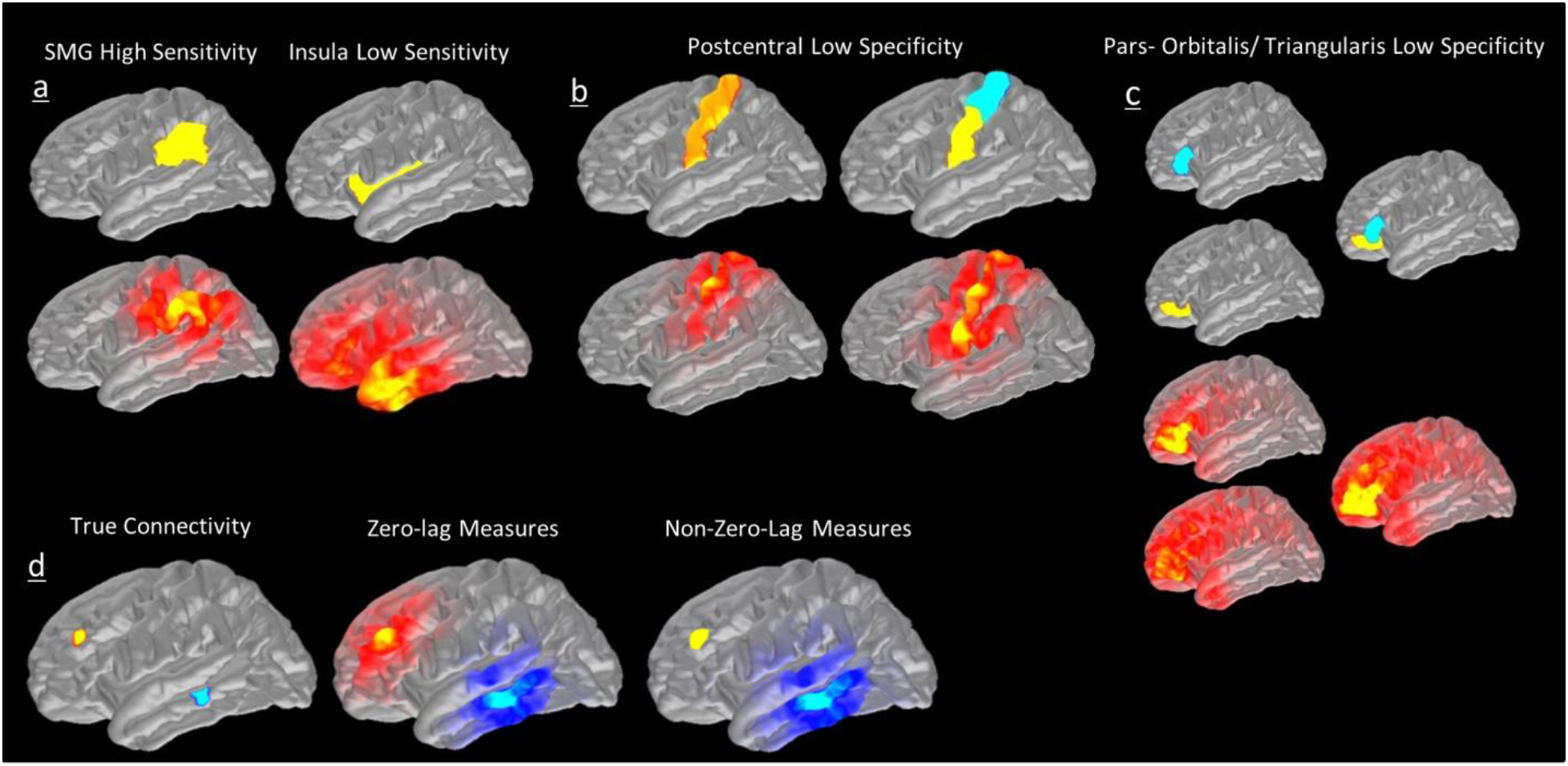
Limitations of the use of anatomical parcellations for EEG/MEG analysis in source space illustrated using CTFs. a) CTFs (bottom) for some parcels (e.g. supramarginal gyrus, left) may peak within the parcel, while for others (e.g. a deep parcel in the insula) the CTF's peak may be at a significant distance from the parcel. b) A single postcentral parcel produces potentially distinguishable CTFs. c) Pars-orbitalis and Pars-triangularis (left; yellow and blue, respectively) are anatomically separate but have largely overlapping CTFs. d) An illustration of how seed-based connectivity is affected by the leakage problem in a hypothetical task where only two fine-grained regions in RMF (seed) and MTG (target) are active and non-zero-lag connected (from theoretical predictions based on CTFs rather than simulation). Left: ideal scenario with no leakage. Middle: in the presence of leakage if a method of connectivity that is sensitive to the zero-lag connections (e.g. coherence) is used. Right: in the presence of leakage if a method of connectivity that is insensitive to the zero-lag connections (imaginary part of coherency) is used.

1. **Sensitivity Problem:** EEG/MEG may not be sensitive to activity from some parcels:
  - While for superficial parcels CTFs may peak within the parcel (e.g. Supramarginal Gyrus, Fig. 1a left), deeper parcels may receive much larger cross-talk from areas close to the sensors than from themselves (e.g. Insula, Fig. 1a right).
2. **Specificity Problem:** Anatomical boundaries might not correspond to the spatial resolution of EEG/MEG:
  a. Large parcels may be split into sub-regions with distinguishable CTFs (e.g. postcentral gyrus, Fig. 1b).
  b. Some distinct anatomical parcels may produce highly similar CTFs, and are therefore indistinguishable from one another due to the limited spatial resolution or EEG/MEG measurements (e.g. Pars Orbitalis and Pars Triangularis, Fig. 1c).

The examples in Fig. 1 also highlight the usefulness of CTFs for the evaluation – and possible construction – of cortical parcellations for EEG/MEG connectivity analysis.

#### 2.1.4 Both zero-lag and non-zero-lag connectivity are affected by leakage

Signal leakage causes activity in one area to be estimated in nearby areas with no time delay; thus there will be zero-lag phase difference between the actual activity and the “leaked” activity (Brookes et al. 2012; Hipp et al. 2012). Therefore, connectivity methods that are insensitive to zero-lag correlations such as phase lag index (PLI) or imaginary part of coherency (ImCOH), have been suggested to overcome the leakage problem to some extent (Stam et al. 2007; Nolte et al. 2004). However, as has been pointed out in some previous studies (Colclough et al. 2015), even though insensitivity to the zero-lag connections can alleviate the problem, non-zero-lag methods are still affected by leakage.

We illustrate the principle of this problem using CTFs in Fig. 1d. Let us consider a case where activity in rostral middle frontal (RMF) cortex and middle temporal gyrus (MTG) show non-zero-lag connectivity. In an ideal scenario with no leakage, the whole-brain seed-based connectivity with seed in the RMF should only produce connectivity with MTG (blue area in the Fig. 1d). However, in a realistic scenario with leakage, two outcomes are possible: 1) If a connectivity measure which is sensitive to zero-lag connections such as Pearson Correlation or Coherence is used, high connectivity will be found between the active sources as well as their leakage domain (Fig. 1d middle); 2) If a non-zero-lag connectivity measure such as imCOH is used, the spurious connectivity between RMF seed and its surrounding areas (i.e. RMF “realm”) will be resolved but results will still be affected by the “blurring” (referred to as inherited connectivity in (Colclough et al. 2015)) around the MTG source (Fig. 1d right). This is due to the fact that the whole neighbourhood of MTG is in non-zero-lag connection to the RMF. It is worth noting that the same argument can be brought for the bivariate directed connectivity methods such as Granger Causality (GC); i.e. if RMF Granger-causes activity in MTG, it will show spurious GC to the neighbourhood of the MTG too. However, generalisation to the multivariate connectivity methods is less straightforward which will be discussed in Appendix A.

## 3 Materials and Methods

### 3.1 EEG/MEG data acquisition and pre-processing

Our results are based on real datasets collected from 17 healthy subjects who participated in an event-related visual word recognition experiment to obtain head-models and noise covariance matrices of pre-stimulus baseline intervals for source estimation. EEG and MEG data were acquired at the MRC Cognition and Brain Sciences Unit, Cambridge, UK, using a Neuromag Vectorview system (Elekta AB, Stockholm, Sweden), which contained 204 planar gradiometers, 102 magnetometers, and a 70-channel EEG cap (EasyCap GmbH, Herrsching, Germany). Individual anatomical T1 MRI scans were acquired using a 3T Siemens Tim Trio scanner at the MRC Cognition and Brain Sciences Unit, using a 3D MPRAGE sequence. A 3Space Isotrak II System (Polhemus, Colchester, Vermont, USA) was used to digitise the positions of 5 Head Position Indicator (HPI) coils that were attached to the EEG cap, 3 anatomical landmark points (left and right ears and nasion), and 50-100 additional points, in order to ensure an accurate co-registration with MRI data. The pre-processing steps for EEG/MEG data (used for the computation of noise covariance matrices) included Neuromag maxfilter (Version 2.0), bad channel interpolation, band-pass filtering between 1-48Hz and ICA for EOG and ECG artefact removals (in our simulations this was relevant for the computation of the noise covariance matrices). MRI preprocessing was performed in the Freesurfer software (Version 5.3; http://surfer.nmr.mgh.harvard.edu/) and EEG/MEG analyses were performed in the MNE python software package (version 0.9) http://martinos.org/mne/stable/mne-python.html). The ICA analysis was performed using FastICA algorithm (Hyvärinen and Oja, 2000) as included in scikit-learn python package (Pedregosa et al. 2011) and implemented in MNE-Python meeg-preprocessing package. As the first step, the dimensionality of the data was reduced using principal component analysis (PCA), by keeping PCs that explain 99% of the data variance and withdrawing the rest. Next, ICs that highly correlated with either of the two ElectroOculoGram (EOG) channels were found by computing Pearson correlation coefficients between each EOG channel and all the IC time courses and converting them to z-score. A maximum of two ICs that showed supra-threshold correlation coefficients (iterative z-score>3) were marked as bad. A maximum of three additional ICs were removed for the ElectroCardioGram (ECG) artefact, however, since there was no ECG recordings in the data, ECG epochs were created from a MEG channel.

### 3.2 Head model and source estimation

Boundary element models (BEMs) were derived from structural MRIs for each subject. Coregistration between MRI and EEG/MEG coordinate systems was achieved on the basis of 50-100 digitised points on the scalp surface, which were matched with the reconstructed scalp surface from the FreeSurfer software. FreeSurfer was used for MRI segmentation and the results were further processed using the MNE software package (Version 2.7.3). The original cortical surface (consisting of more than 160,000 vertices) was down-sampled to a tessellated grid where the average edge of each triangle was approximately 2.5mm, resulting in 20484 vertices in the downsampled cortex (Segonne et al. 2004). A three-layer BEM consisting of 5120 triangles per layer was created from combined EEG/MEG from scalp, outer skull surface and inner skull surface respectively. The noise covariance matrices for each dataset were computed and regularised in a single framework which computes the covariance using empirical, diagonal and shrinkage techniques and selects the best fitting model by log-likelihood and three-fold cross-validation on unseen data (Engemann & Gramfort 2015). Baseline intervals of 500 ms duration pre-stimulus were used for the estimation of noise covariance matrices. The resulting regularised noise covariance matrices were used to assemble the inverse operators for each subject using an L2 minimum norm (MNE) estimator with loose orientation constraint 0.2 (Lin et al. 2004) and no depth weighting.

### 3.3 EEG/MEG-adaptive parcellations

As outlined in the introduction, we aim to obtain a parcellation of the cortical surface into parcels that, according to their CTFs, are sensitive to activity originating from or around them, but are relatively insensitive to leakage from other parcels. In the first approach, we started from existing standard anatomical parcellations, and optimised them using a modified split and merge (SaM) algorithm. In the second approach, we started with no prior parcellation and created an optimal set of parcels using a region growing (RG) algorithm. Both SaM and RG belong to the so-called region-based family of image segmentation with relatively simple and robust implementations of algorithms (Gonzalez & Woods 2007). These algorithms were preferred over more complex and less frequently used methods and also over edge-based family as another simple and common family of segmentation algorithms (Pham et al. 2000; Dutta et al. 2016). The latter was due to the fact that edge-based algorithms aim to form contours around the distinguishable parts of an image by setting some criteria for edge detection (e.g. gradient). Therefore, clear boundaries are typically required and the algorithms are sensitive to the presence of noise (Pal & Pal 1993). Hence, since CTFs of different brain areas are not necessarily clearly separated and noise levels in the data can be high, region-based methods were preferred. Moreover, among the region-based algorithms, those that require a pre-specified number of regions (segments) such as clustering methods (Pham et al. 2000) were not suitable. Instead, one of the main purposes of the current study is to recruit algorithms that yield the optimal number of parcels in the brain. Both SaM and RG are simple, fast and robust against noise and can be expected to yield coherent focal regions in the brain.

A flowchart of different steps of analyses is shown in Fig. 2.

**Figure 2.**
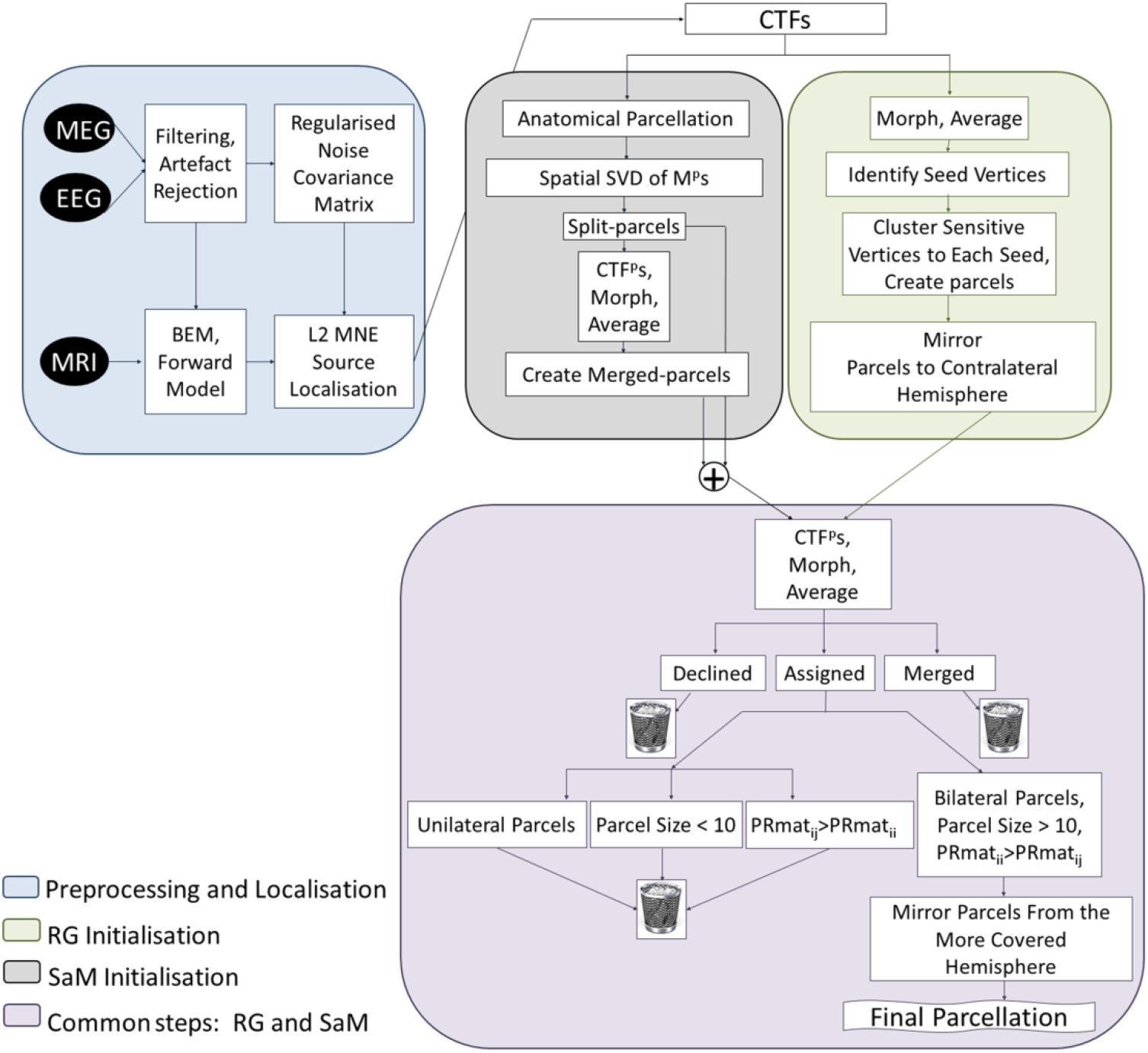
A flowchart of data analysis and parcellation algorithms. Preprocessing and localisation steps (blue box) can change depending on the study and CTFs and subsequent steps will change and adapt accordingly. M^p^: a subset of R matrix corresponding to each parcel, CTF^p^: CTFs of each parcel at all the brain vertices, PRmat: parcel resolution matrix, RG: Region Growing, SaM: Split and Merge.

#### 3.3.1 Leakage and Parcel Resolution Matrices (PRmat)

The starting point for our algorithms is the Parcel Resolution Matrix (PRmat). While the resolution matrix R (Equation 2) describes cross-talk among all vertices, PRmat describes normalised cross-talk among parcels. Below, we will describe the computation of PRmats. Let us assume that at one stage within our algorithms, we have N_parcel_ parcels with N_v_ overall vertices and N_p_ vertices per parcel p.

- First, we compute absolute values of CTFs at each vertex, as we are only interested in the amount of leakage. We will still refer to these as CTFs for simplicity.
- We arrange all CTFs for vertices within a parcel p as rows of a matrix M^p^. Thus, M^p^ is a submatrix of R, containing only those rows corresponding to vertices in the parcel p.
- We compute the singular-value decomposition (SVD) along rows of all matrices M^p^. We then represent each parcel by the first eigenvector CTF^p^ along rows (i.e. across CTFs).
- Second, we define **PRmat**, where each element PRmat_ij_ describes leakage from parcel i to parcel j, normalised by the amount of leakage it receives from all parcels:

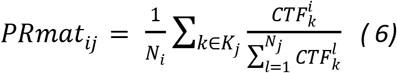

where K_j_ refers to the set of indices for vertices in parcel j and 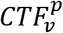 is the cross-talk of parcel p at vertex v. The normalisation ensures that values in PRmat are between 0 and 1. The ideal PRmat is an identity matrix, and our purpose is to obtain parcellations for which the similarities between the actual and an ideal PRmat are maximised.

#### 3.3.2 A CTF and neuro-anatomy based split-and-merge (SaM) segmentation algorithm

In this section we examined Desikan-Killiany (DKA, 68 parcels) and Destrieux (DA, 148 parcels) Atlases that are defined in the fsaverage space in the Freesurfer software. Two different parcellations were used since DKA provides a coarser sampling of the cortex while DA provides a more fine-grained sampling of the cortex. This enabled us to observe the effect of the initial parcel sizes on the final results of parcellation modifications. We modified the parcellations using a CTF-informed algorithm similar to the split-and-merge algorithm in digital image processing literature (Haralick & Shapiro. 1985; Gonzalez & Woods 2007). Split-and-merge algorithm which is one of the established methods for image segmentation typically starts from a whole image and utilises an iterative process to divide the image into as many “homogeneous” segments as possible. The homogeneity is defined based on the image properties, for example, one implementation of the algorithm might seek to segment an image based on constant standard deviation inside each segment. If the homogeneity criterion is not satisfied inside a segment, that segment will be split into several equal-sized sub-segments and the homogeneity criterion will be checked inside each of these new segments. This procedure is iterated until no further splitting is possible. At this point, the algorithm searches for the segments that might have been over-split during the splitting and merges them together. To this end, the segments that show similar properties based on some predefined criterion (e.g. pixel colour or intensity) will be merged in an iterative procedure until no more merging is possible.

Here, we have adapted a similar idea and have defined the split, merge and homogeneity criteria based on CTFs. On the one hand, parcels that are too large to be represented by one CTF should be split up. On the other hand, if CTFs of two parcels overlap substantially, those parcels cannot be distinguished using EEG/MEG (Fig. 1b, c). Furthermore, if EEG/MEG is not sensitive to activity from a parcel, it should be omitted from the parcellation (Fig. 1a). Therefore, CTFs and resolution matrices can be used to inform the splitting and merging in order to parcellate the cortex into the optimal number of distinguishable parcels. As will be elaborated in the next subsections, the SaM algorithm used in this study is a non-iterative version of the original SaM algorithm described above.

##### 3.3.2.1 Splitting criterion

The purpose of the first step – splitting – was to identify large parcels (e.g. Fig. 1b) and split them into several sub-parcels. For a particular parcel,

- We determined the number of principal components (PCs), NPC, needed to explain more than 90% across their CTFs (determined from an SVD of matrix M^p^ in 3.3.1).
- If N_PC_> 1, we split the parcel into N_PC_ sub-parcels along its longest spatial axis. This is done by finding the principal eigen-axis of the label on the spherical surface, projecting all the coordinates of the label vertices on this axis, and dividing them at equal intervals.

In order to obtain a fixed number of sub-parcels across hemispheres per subject as well as across all subjects in the data set, we added the following constraints:

- In order to obtain consistency across hemispheres, the minimum of N_PC_ for the corresponding parcels in the left and right hemispheres was assigned to both parcels. We consider the oversplitting of parcels, i.e. multiple parcels that contain the same information, as less desirable than under-splitting, i.e. a parcel that potentially covers a larger area than necessary.
- In order to obtain one splitting number for each parcel across subjects, we looked at the distribution of N_PC_ across subjects and assigned the mode (i.e. the most frequency number, the minimum number if multiple modes) of this distribution to the parcel.

This resulted in a “split-parcel” parcellation, which was used for further processing.

##### 3.3.2.2 Homogeneity criterion

After creating a parcellation consisting of split-parcels, we tested for all individual vertices whether we could reassign them to one of the new parcels, whether we should drop them because no new parcel was sensitive to them, or whether they were candidates for a later merging procedure. For this purpose, CTF^p^s for the split parcels were computed and morphed to the average brain, in order to be averaged over subjects. Thereafter, we assigned each of the vertices in the average brain to a maximum of one split-parcel. A vertex was assigned to a split-parcel only if it was

1. Sensitive to that split-parcel (**sensitivity**)
2. Significantly more sensitive to that split-parcel compared to all other split-parcels in the brain (**specificity**).

Sensitivity and Specificity were defined as follows.

**Sensitivity**: We removed the vertices that were not sensitive to any split-parcels. To this aim, for every vertex, we tested for every split-parcel whether the split-parcel's CTF^p^ value at this vertex was equal or more than half of the maximum of the split-parcel's CTF^p^ values anywhere in the brain. If this was the case, that vertex was considered sensitive to that split-parcel. Vertices that were not sensitive to any split-parcels in the brain were removed from further analysis.

**Specificity**: For every pair (i,v) of split-parcel i and vertex v, we quantified the relative cross-talk that vertex v receives from split-parcel i compared to all other N split-parcels as the z-score Z_iv_:

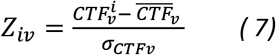

where 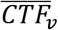 denotes the average of CTF^p^ values of split-parcels at vertex v, and σ_*CTF*v_ denotes the standard deviation across CTF^p^ values from all split-parcels at vertex v, respectively.

Based on these z-scores, we classified vertices into one of three categories:

a. Declined vertices: If no split-parcels showed a z-score above 3 for a vertex, it indicated that the vertex was not specifically sensitive to the CTF^p^ s of any of the split-parcels in the brain. These vertices were removed from further analysis.
b. Assigned vertices: Using a winner-takes-all approach, if the split-parcel with the highest z-score for a vertex had a z-score above 3 and at least 1 standard deviation higher than the runner-up, that vertex was assigned to this split-parcel.
c. Merge candidate vertices: vertices that showed similarly high sensitivity to two split-parcels were marked for the merging procedure (see sub-section 3.3.2.3 below). In other words, merge candidates consisted of a vertex (or a patch of vertices) that showed high z-score (>3) to a pair of split-parcels but the difference between the z-scores was less than 1.

##### 3.3.2.3 Merging criterion

The aim of the third step was to create a set of merged-parcels based on the “merge candidate vertices” described above. For this purpose, for each pair of split-parcels in the brain, a group of vertices that were similarly highly sensitive to those split-parcels were clustered together as a new “merged-parcel”. These merged-parcels resulted from two scenarios:

1. If two original split-parcels were too finely separated and not distinguishable using EEG/MEG (e.g. parcels in Fig. 1c), they were completely merged together.
2. If CTF^p^s of two split-parcels were partially overlapping, a third region might have emerged from that overlapping region.

Of these new merged-parcels, those that were of equal-size or larger than the smallest original split-parcel in the brain were kept for further analysis.

As an example, vertices that were equally sensitive to both superior temporal and middle temporal gyri were clustered as a new merged-parcel called superior-temporal_middle-temporal.

##### 3.3.2.4 Final homogeneity evaluation

The procedure above resulted in a new parcellation (consisting of the original split-parcels and the new merged-parcels shown in Fig. C. 1 in Appendix C), based on splitting, merging and homogeneity criteria. However, these criteria used CTF^p^s based on the initial parcellation. We therefore need to optimise the new parcellation based on its own PRmat.

Therefore:

- Step 3.3.2.2 (homogeneity criterion) was repeated for the modified list of split- and merged-parcels.Those parcels that could win at least 10 vertices were kept and the rest of the parcels were declined.
- The **PRmat** was computed for the modified parcels and if any off-diagonal elements of a particular parcel were higher than the diagonal element, that parcel was removed.
- In order to obtain a consistent parcellation across hemispheres, those parcels that survived the above criteria in only one hemisphere were removed. Moreover, in order to obtain a symmetrical parcellation in the end, parcels were kept in the hemisphere that showed more coverage and mirrored to the opposite hemisphere.

#### 3.3.3 A CTF-based region growing segmentation algorithm for the parcellation

Region growing is another algorithm of image segmentation which typically starts by randomly selecting a voxel (pixel) as the first “seed” in an image. Then, based on a pre-specified similarity criterion (e.g. colour or intensity), neighbouring voxels are grouped together with the seed voxel, leading to a growing region around the seed until no more voxels can satisfy the similarity criterion to connect to the cluster (Gonzalez & Woods 2007). Thereafter, a new seed outside the existing cluster is randomly selected in the image and the same procedure is iterated until all the voxels in the image are assigned to one cluster. In this section, we have adopted a similar idea and have used CTFs to define the similarity criterion to grow regions around the vertices in order to create and modify parcels in the brain. Therefore, we started the parcellation at the single-vertex level with no prior parcels and created parcels using the following steps:

##### 3.3.3.1 Finding seed vertices

The main purpose of the first step was to identify the “seed vertices”, i.e. vertices that show high sensitivity based on the CTFs. Therefore:

- The resolution matrix (**R**) was computed for all vertices (section 2.1.2 with rows representing CTFs at each vertex.
- Sensitivity and specificity steps described in section 3.3.2.2 were applied to the rows of the resolution matrix in order to find the sensitivity of each vertex to leakage from all other vertices. In other words, every vertex was treated like a “split-parcel” in 3.3.2.2, and then we tested (i.e. using a winner takes all approach with significantly highest z-score > 3) whether other vertices will be grouped with each vertex. Those vertices that could “win” more than one vertex were marked as seeds.

##### 3.3.3.2 Growing regions surrounding the seeds

The second step comprised of growing regions around the seeds. For this purpose, we sorted the seeds in a descending order with the first seed being the “strongest” and created regions in succession following this order.

- Seeds were sorted based on their sensitivity to themselves; i.e. the strongest seed (seed 1) had the highest z-score for itself (section 3.3.2.2).
- All vertices that showed sensitivity to seed 1, i.e. produced higher cross-talk in seed 1 than the half maximum of the CTF values of this seed, were clustered together as parcel1.
- In an iterative procedure, parcel_n+1_ was created from the vertices outside all parcel_i_ with i <= n, with the same half maximum criterion.
- To obtain an inter-hemispheric symmetry of the parcels, the created parcels of the hemisphere with more winner seeds were mirrored to the opposite hemisphere using MNI coordinates. These created parcels are shown in Fig. C. 1 in Appendix C.

##### 3.3.3.3 Modifying the parcels

- The same procedures as those described in 3.3.2 (except for the splitting step) were applied to the parcels created by the region-growing (RG) algorithm to obtain the final RG parcellation.

#### 3.3.4 Parcellation performance metrics

We used PRmats to evaluate the performance of different original and modified parcellations. As explained earlier, the PRmat is computed by finding the normalised CTF values produced by each parcel at the location of all other parcels. If a parcellation consists of fully distinguishable parcels, the PRmat should be an identity matrix. Here we introduce two metrics to evaluate a parcellation's performance:

- The **Sensitivity Index** (S_ind_) measures the sensitivity of parcels to themselves by taking the mean of the diagonal elements of the PRmat.

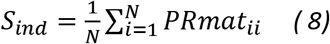

where N is the number of parcels in the parcellation. The ideal value would be 1.
- The **Distinguishability Index** (D_ind_) is the correlation between the actual **PRmat** and the identity matrix of the same dimensions.

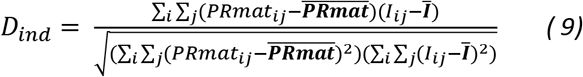 Where ^—^ denotes the average of matrix elements and **I** is the identity matrix.

Furthermore, we computed the rank and condition numbers of PRmats to make comparisons between the original anatomical and modified parcellations. This rank of the resolution matrix is far lower than the ideal rank, i.e. the number of vertices. This means that the rows of R, i.e. the CTFs for all vertices, are not linearly independent, and activity in all vertices cannot be estimated independently of each other. Therefore, the condition number of this matrix will be infinity. This is different for PRmat, where the parcels were chosen to minimize overlap of CTF^p^s. A low condition number (especially around the value 1) would indicate that all CTF^p^s are non-overlapping, and that an inversion of PRmat (e.g. for leakage correction) would be stable.

Hence the number of degrees of freedom is smaller than the number of rows/columns. Considering that PRmat is scaled between 0 and 1, we computed the rank with a heuristic tolerance of 0.05. It is worth noting that this value is much higher that the numerical precision for rank computation, however, it shows that if similarities between one row of the PRmat and a linear combination of all other rows are higher than 95%, that row will not be considered as independent from other rows. A high condition number is indicative of an ill-conditioned parcel resolution matrix, i.e. the estimated sources (output) can be very sensitive to small changes in the actual sources (input). A high condition number indicates that if the PRmat was to be inverted (e.g. to perform leakage correction based on the final PRmat) the results will be unreliable.

Additionally, for each parcellation we computed the coverage which is the total number of vertices that are included in the parcellation.

### 3.4 Simulation with realistic levels of noise

In this section, we compare the performance of different parcellations for network reconstruction using simulated data with realistic levels of noise. For this purpose, we simulate several hundred realistic datasets in order to evaluate the performance of different anatomical and modified parcellations. To our knowledge, this provides the first comparison of the effect of the choice of various parcellations on reconstruction of realistically simulated EEG/MEG networks. Our simulations are based on head models, forward and inverse operators of the 17 subjects described in 3.1 and 3.3. We use coherence as the measure of connectivity (edge strength) to reconstruct the simulated networks based on each of the anatomical and modified parcellations. Finally, we estimate the significant connections in each network and compare the results to the simulated ground truths. Details of each step are outlined in the next subsections and a flowchart of different steps is depicted in Fig. 3. All simulations were carried out in python, and where appropriate (e.g. forward and inverse modelling), we used the mne-python software package.

**Figure 3.**
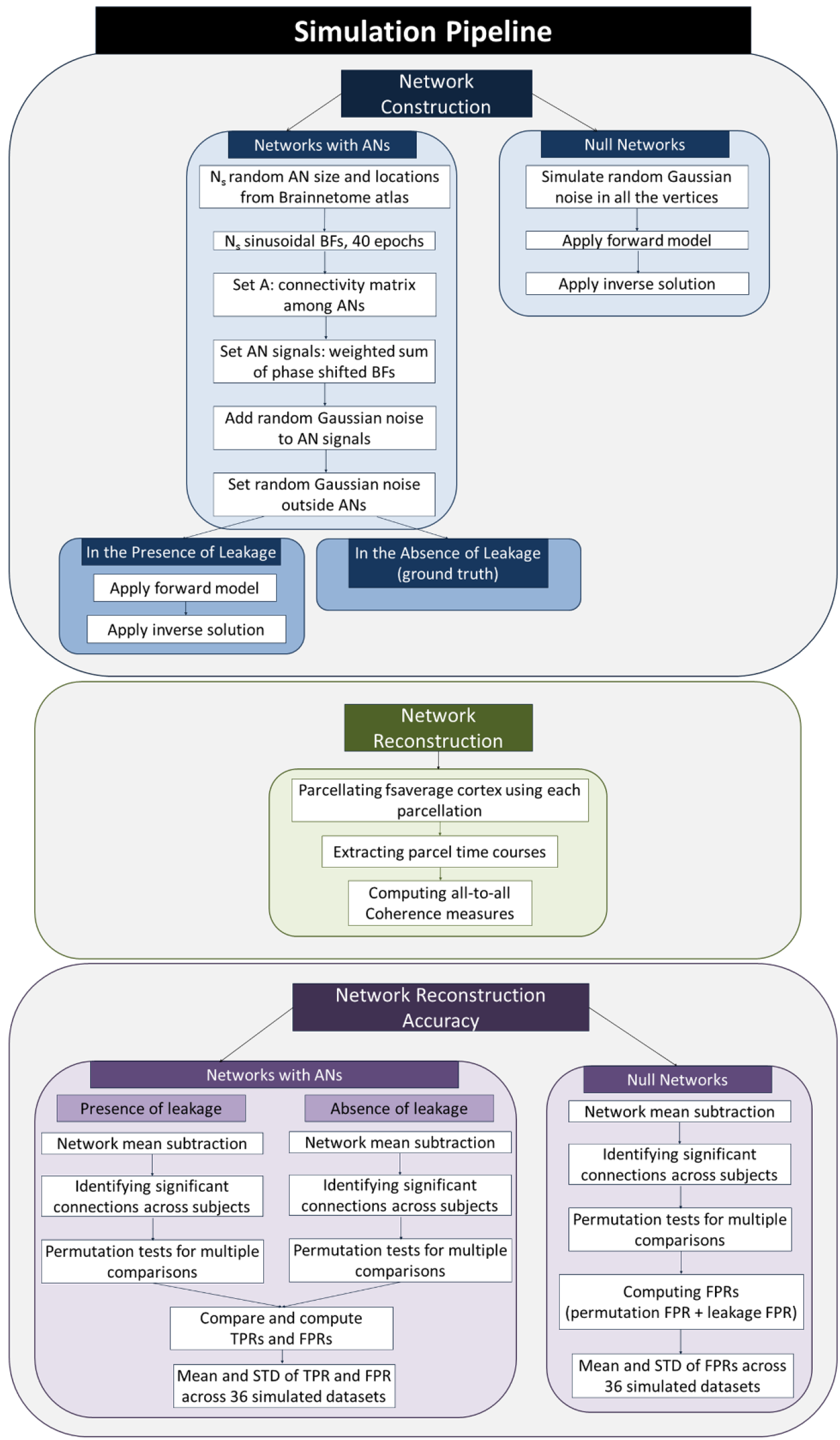
Flowchart of the simulation pipeline consisting of three main steps of network construction, network reconstruction and network reconstruction accuracy. AN: Active node, BF: Basis function, TPR: True positive rate, FPR: False positive rate, STD: standard deviation.

#### 3.4.1 Network Construction

We simulated a range of networks by varying the number of active seeds (3, 5, 10, 15) in the brain, the percentage of connections among those seeds (25%, 50% and 100%) and SNR of the data (1.0 and 3.0). For each of these cases, 36 random datasets were created in two scenarios: absence of leakage (reference ground truth) and presence of leakage. Additionally, we simulated 36 null networks (i.e. random networks with no significant connectivity patterns). These simulations yielded 1764 datasets overall, each consisting of 17 subjects.

##### 3.4.1.1 Location and size of active sources

Each network was initiated by randomly selecting N_s_ seeds on the cortex (fsaverage space). We defined the active seeds by randomly drawing parcels from the cortical areas of the Brainnetome functional atlas of the brain ((Fan et al. 2016) Fig. B. 1a in Appendix B). This approach of defining the source locations has two main advantages: a) defining seeds based on a canonical functional atlas provides a realistic representation of size and location of likely functional seeds in the brain; b) the size and locations of seeds are independent of the choice of parcellations that will later be used for network reconstruction. This prevents biases towards any of the parcellation approaches. It is worth noting that some previous studies have tested their parcellations using sources that corresponded to active parcels in their parcellation (e.g. (Korhonen et al. 2014)) so as to obtain a one-to-one correspondence between active seeds and cortical parcels. However, in comparison between different parcellations with different number and locations of the parcels, which is the case in the current study, that approach would result in a bias in favour of one parcellation or the other.

##### 3.4.1.2 Simulated signals and connectivity patterns

N_s_ sinusoidal signals for 40 epochs (duration 725ms including 125ms baseline) were simulated in the randomly selected seed locations. All of the vertices within each active node (AN) were assigned the same signal and the rest of the vertices in the brain were assigned noise. In order to systematically vary connectivity in our simulated networks, we created activation time courses at each AN as a weighted sum of a fixed set of basis functions:

a. a) As basis functions (BFs), we first simulated N_s_ signals, each across all 40 epochs. These BFs were arranged as rows 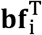 of a matrix **M_BF_** (size N_s_⨯N_t_, N_t_: number of samples across epochs, Equation 10), each defined based on a function *f_i_*(*t*).
b. b) The frequency of each BF was randomly selected from a list of frequencies obtained by dividing the interval of 10-40Hz into N_s_ equally-spaced frequencies. The phase of each BF randomly varied across the epochs in order to ensure no coherence between each pair of BFs.
c. c) We then computed the activation time courses at each AN (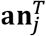, Equation 11, defined by functions *g_i_*(*t*)) as a weighted sum of phase-shifted versions of these basis functions. Each AN was given a randomly selected phase (φ_*j*_, Equation 13) which remained constant over epochs. Therefore, if two ANs share the same BFs, there will be significant connectivity between them. Then, the signals for all ANs were arranged in a matrix **M**_AN_ (size N_s_⨯N_t_, N_t_: number of samples across epochs, Equation 11), such:

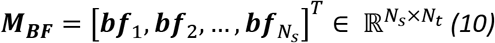

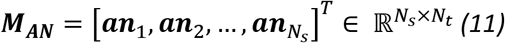

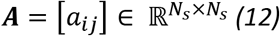

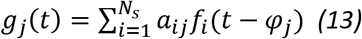 **M**_BF_ is the matrix of BF signals of size N_s_⨯ N_t_, is the phase shifts for the j^th^ AN and A is the desired connectivity matrix:
d. In the Equations 12 and 13, connectivity among ANs was defined by a binary matrix **A** which is of size N_s_ x Ns. Each row corresponds to one AN, and determines the contribution of each BF to its activation time course. Therefore, the BFs that are given ones in the row of **A** that corresponds to each AN will contribute to that specific AN. We imposed the following constraints on matrix **A**:
  i. The diagonal elements of **A** were all set to one, thus, each BF is inherent to one of the ANs. This ensures that all ANs are active (i.e. none is flat or all-noise) even if all the off diagonal elements of a row of **A** are zero.
  ii. Since we introduced random non-zero phase-shifts between nodes, the resulting signals will have non-zero-lag connectivity.
  iii. For each network scenario, 25%, 50% or 100% of all possible connections among the ANs were set to be non-zero. These included the connections obtained by setting the corresponding elements of **A** matrix to one and connectivity through connections to a common third source. The latter is taken into account since if, for example, nodes AN_1_ and AN_2_ are both connected to AN_3_ through sharing BF_3_, the network reconstruction (see section ‎3.4.2 below) will reveal a significant connectivity between AN_1_ and AN_2_ even if the corresponding element of **A** matrix is not set to one. It is worth noting that taking indirect connections into account is not conventional since matrix **A** fully describes the connectivity patterns of the “constructed networks”. However indirect connections will become important at the level of “network reconstruction” since our functional connectivity metrics cannot distinguish between direct connections and indirect connections through a third source (multivariate metrics might alleviate this problem but cannot solve it). Thus, if indirect connections are not taken into account when constructing the network, reconstruction of a network with e.g. 10 seeds and 50% of connections might look very similar to the reconstruction of a network with 10 seeds and 100% of connections and hence, comparisons between some percentages of connectivity among seeds becomes trivial. This is more important for higher number of active nodes (e.g. 10 and 15 seeds compared to 3 and 5).
  iv. The relative phases of ANs, the elements of A matrix and the frequencies of each BF are selected randomly, hence, the values of coherence among the ANs vary between 0 and 1 depending on the noise level and number of connections, which we assume to be the case for realistic brain networks.
  After simulating the sinusoidal signals of the ANs, we added noise to them. Furthermore, we simulated random noise in all the vertices outside the active source locations. These networks were constructed with two levels of SNR: 1 and 3 in order to explore the effect of noise on the parcellation performances. SNR was defined as the square root of mean square signal (i.e. after 0ms) divided by the standard deviation of noise.

Each network was constructed in two scenarios:

- In the presence of leakage: Sources were simulated in the fsaverage source space, morphed to the single subject source space, projected onto the sensor space using the individual forward models and projected back to the source space using the individual inverse operators (described in 3.3). Each vertex time course was extracted from the source component normal to the surface, and the obtained activation maps were morphed back to the fsaverage source space.
- In the absence of leakage (reference ground truth): The simulated sources in the brain were analysed directly, without the application of forward and inverse operators. This will serve as the ground truth against which the performance of parcellations will be compared.

##### 3.4.1.3 Null Networks

In addition to the networks elaborated above, we constructed a set of null networks in order to study the performance of parcellations in the absence of true brain connectivity. For this purpose, we simulated noise in every vertex of the fsaverage brain. The simulated signals were morphed to individual head spaces, forward and inverse models were applied to these noise-induced networks in order to obtain leakage-induced networks. And finally, these leakage-induced networks were morphed back to the fsaverage space. Similar to the realistic networks with active nodes, 36 datasets were created from 17 subjects for each of the leakage-induced networks.

#### 3.4.2 Network reconstruction

We used Magnitude-Squared Coherence (COH) and imaginary part of Coherency (imCOH) as two measures of connectivity to reconstruct the simulated networks and compare the performance of different parcellation methods for whole-brain network reconstruction. COH and imCOH are spectral measures of connectivity which can detect both amplitude and phase couplings (Greenblatt et al. 2012; Bastos & Schoffelen 2016). COH is sensitive to zero-lag connections while imCOH is not (Nolte et al. 2004; Bastos et al. 2012). We used imCOH as well as COH to evaluate the consequences of the theoretical issue discussed in 3.1.2; i.e., whether EEG/MEG-adaptive parcellations are useful both for zero- and non-zero-lag connectivity measures. In order to reconstruct each network using these measures:

- We simulated signals in each of the scenarios outlined above which resulted in N_v_ ⨯ t matrix of vertex time courses where N_v_ is the number of vertices in the brain and t is time. As the first step of reconstruction, we parcellated the fsaverage cortex using each of the anatomical and modified parcellations. It is worth noting that each active node (i.e. each parcel of the Brainnetome atlas) contributes to all the parcels in the anatomical/adaptive parcellations that overlap with that source, depending on the number of spatially overlapping vertices. Therefore the extracted time course for each parcel will be determined by the signal of the ANs that it overlaps with plus noise vertices inside that parcel.
- Next, we collapsed the matrix of vertex time courses to a matrix of parcel time courses, TC, of size N x t where N is the number of parcels and t is time. In order to extract the parcel time courses, we used a mean-flipped approach. This approach computes a parcel time course by taking the average of the sign-flipped signals of the vertices within that parcel. The flipping sign is determined based on the source orientation at each vertex within the parcel, with positive indicating outward-flowing currents.
- Thereafter, we computed COH and imCOH on TC and obtained an N ⨯ N connectivity matrix M_con_. COH and imCOH were computed using a multitaper approach with adaptive weights in a broad band frequency of 8-55Hz.

##### 3.4.2.1 Lower coverage of the cortex by the modified parcellations

In the steps described above, the “ground truth” of each parcellation is determined based on that specific parcellation in the absence of leakage. However, it is worth noting that, unlike the anatomical parcellations that cover the cortex fully, the modified parcellations provide only a sparse sampling, so it is likely that some of the randomly selected seed locations do not coincide with any of the parcels (Fig. B. 1) and therefore they will be absent in the “ideal ground truth” as well as realistic scenarios in the presence of leakage. Therefore, we additionally recorded the number of connections in the **A** matrix that were missed due to no coverage of the corresponding ANs using each of the modified parcellations. This will be taken into account in the computation of true positive rates below.

#### 3.4.3 Network reconstruction accuracy

We used statistical analysis in order to evaluate the accuracy of network reconstruction based on each of the parcellations:

- Firstly, for each network, the average value of the absolute values of all connections within that network was used as the baseline and was subtracted from the absolute value of all the elements of the **N ⨯ N** connectivity matrix, **M_con_**. Baseline correction was applied in order to obtain connectivity values that are distributed around zero and are suitable for statistical analysis. Therefore, the elements of connectivity matrix that are below average in some subjects and above average in other subjects are likely due to noise while the connectivity values that are consistently above average are unlikely to be merely due to the noise. Furthermore, the absolute values (relevant for ImCOH) were used because, regardless of the sign of connectivity between two areas, the strength of connections is important for evaluation of statistical significance. It is worth noting that the choice of threshold is often arbitrary and should ideally be tested for a range of different values (Rubinov & Sporns 2010). However, in this study, since we are using average thresholds for reconstructed networks in the presence and absence of leakage (ground truth) in order to make comparisons between the two, testing various thresholds is not strictly required. Furthermore, since the same procedure is applied to both adaptive and anatomical parcellations, we expect no bias in favour of any of the parcellations due to the thresholding.
- For each ground truth network in the absence of leakage, significant connections were identified using one-tailed permutation tests (i.e. only connections that are significantly higher than the baseline are of interest), which included correction for multiple comparisons across connections. These calculations yielded “true significant connectivity” among the parcels in each parcellation.
- Baseline correction and permutation tests were also applied to each realistic network, and then compared against the true connectivity matrix, with two groups of connections identified:
  - True positives: Significant connections that were identified accurately in the realistic networks divided by the overall number of true connections. Note that we included the number of missed connections due to no coverage of some ANs by modified parcellations (see 3.4.2.1) in calculation of true connections.
  - False positives: non-existent connections in the ideal scenario that were incorrectly marked as significant in the realistic networks divided by the overall number of zero connections in the ground truth.
  These metrics were computed for each random dataset and averaged across 36 iterations. All of the evaluation steps were applied to the results of connectivity from both COH and imCOH.

## 4 Results

### 4.1 Parcellation results

#### 4.1.1 Split-and-Merge algorithm (SaM)

We tested the split-and-merge (SaM) algorithm (section 3.3.2) on two standard anatomical parcellations in Freesurfer: Desikan-Killiany and Destrieux Atlases that are shown in Figure 4a, c with the corresponding Parcel Resolution Matrices (PRmat: relative between-parcel leakage values, see 3.3.1) shown in Figure 4b, d, respectively.

**Figure 4:**
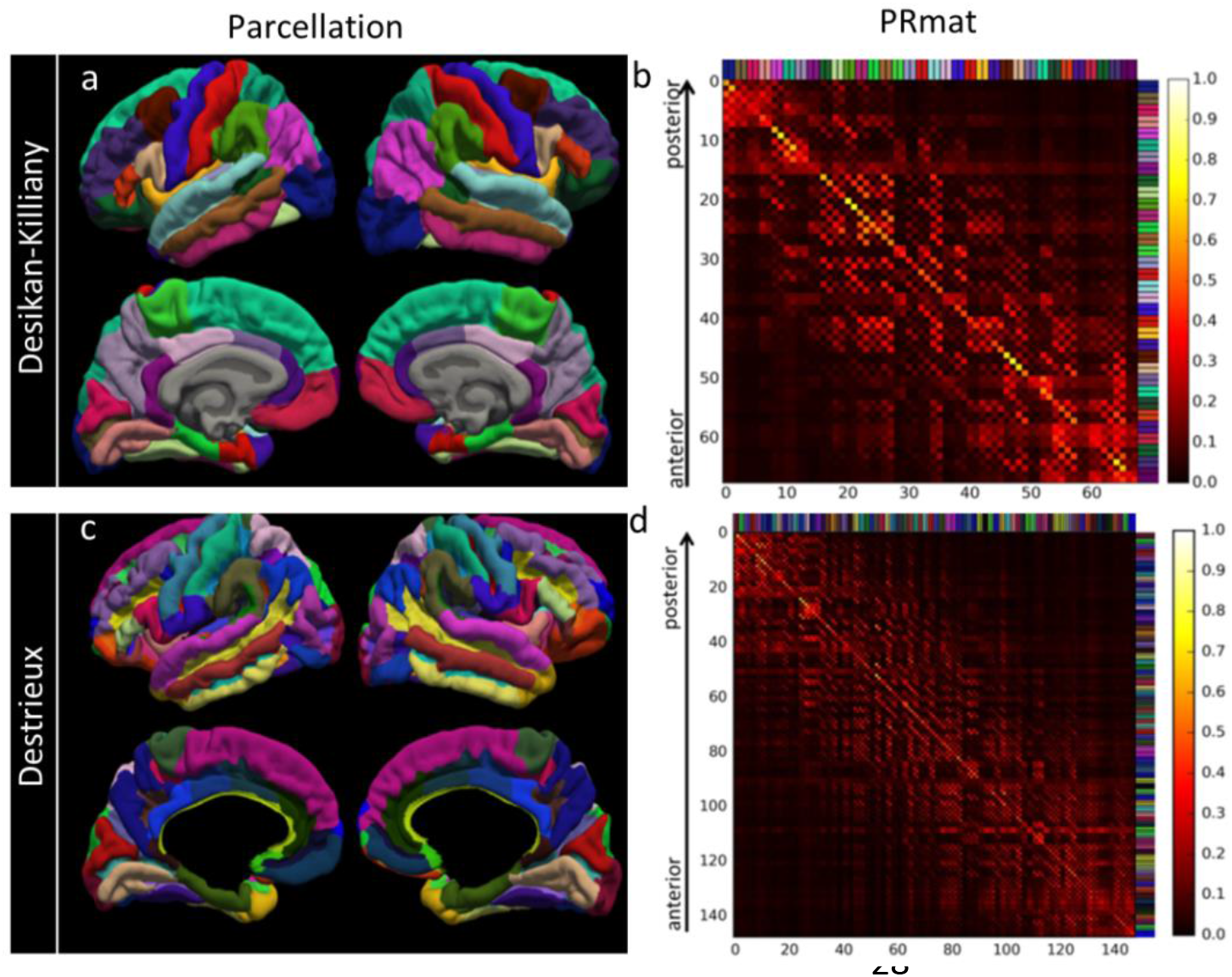
a) The original anatomical Desikan-Killiany Atlas with 68 parcels bilaterally; b) P arcel Resolution Matrix (PRmat) of Desikan-Killiany Atlas. Rows s how average normalised CTF of each parcel (equation 6). c) The original anatomical Destrieux Atlas with 146 parcels bilaterally; b) PRmat of the parcellation. C olour labels along rows and columns of the PRmats correspond to those used for the parcellations.

##### 4.1.1.1 Desikan-Killiany Atlas

The original Desikan-Killiany Atlas included 68 parcels with sensitivity index S_ind_ of 0.47 (i.e. the leakage value that each parcel received from itself relative to the rest of the parcels in the brain) and distinguishability D_ind_ of 0.50 (i.e. correlation between the PRmat and an ideal identity matrix) (Table 1). The SaM algorithm resulted in 316 parcels at the intermediate step (Fig. C. 1a, b; Appendix C), from which 74 regions survived to the final parcellation that is shown in Figure 5a together with the corresponding PRmat. Compared to the original parcellation, S_ind_ and D_ind_ increased by 38% and 22% and reached 0.65 and 0.61 respectively (Table 1) and provided a sparser sampling of the cortex including 4079 vertices.

**Table 1.**
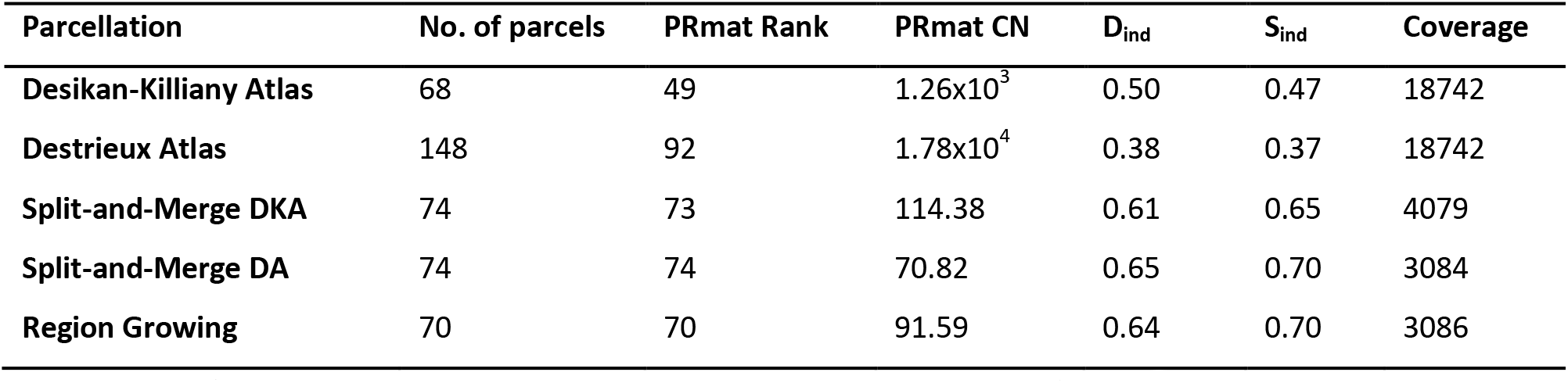
A summary of the performance of the original and modified parcellations. D_ind_: Distinguishability index, S_ind_: Sensitivity index, PRmat: Parcel Resolution Matrix, CN: Condition number

##### 4.1.1.2 Destrieux Atlas

The original Destrieux Atlas consists of 148 parcels and is shown in Figure 4c with PRmat in Fig. 4d. In comparison to the Desikan-Killiany parcellation, the PRmat of this parcellation shows less similarity with an identity matrix, indicating a more blurred estimation of activity for each of the parcels (Table 1). This difference suggests that the original Desikan-Killiany is a better match to the EEG/MEG spatial resolution than Destrieux. S_ind_ and D_ind_ of Destrieux Atlas were 0.37 and 0.38, respectively, and improved to 0.7 and 0.65 for the 74 parcels that survived the parcellation modification, providing an 89% and 71% improvement in these indices, respectively. The parcellation covered 3084 vertices of the cortical surface. The intermediate and final parcellation/PRmat for the modified Destrieux Atlas are shown in Fig. C. 1 c, d and Fig. 5b respectively. Comparison to Fig. 4d, as reflected in increased S_ind_ and D_ind_ values above, shows a clear improvement. Note that in Fig. 5b, parcels that showed maximum overlap with each of the modified parcels from the Desikan-Killiany are colour-matched to Fig. 5a for visual comparison.

**Figure 5.**
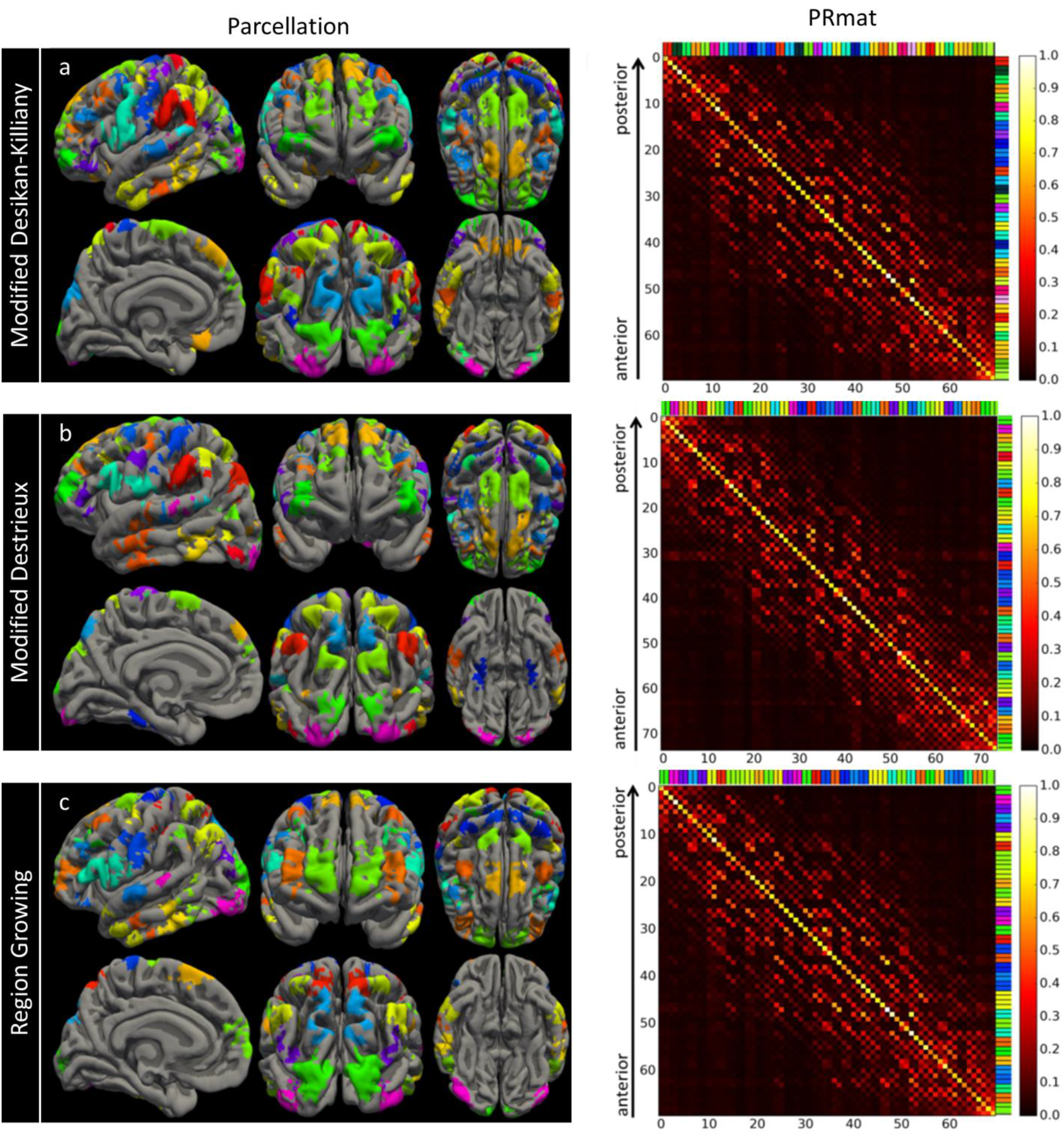
Final parcellations (left) and PRmats (right) for a) SaM algorithm based on Desikan-Killiany Atlas; b) Destrieux Atlas; c) Region growing algorithm.

Despite having twice the number of initial parcels, the SaM algorithm converged at 74 parcels for both atlases. This can be considered as an indicator of the robustness of the parcellation algorithms against the initial choice of parcellation.

#### 4.1.2 Region Growing algorithm (RG)

The Region Growing Algorithm does not require an anatomical parcellation as a starting point, but creates a parcellation based on the resolution properties of all the vertices. The first step of RG algorithm identified 174 seed vertices (Fig. C. 1e) in the left hemisphere and parcels were grown surrounding each of these seeds using the criteria described in 3.3.3. The split and merge criteria were applied to these created parcels and resulted in a 70-parcel parcellation with S_ind_ of 0.7, D_ind_ of 0.64 and a sparse sampling of the whole cortex, covering 3086 out of 20484 vertices in the brain (Table 1). The final parcellation showed notable similarities and differences to the parcellation modification of the anatomical atlases (Fig. 5c). A direct comparison of the overlaps and differences of the final parcellations are conducted in section 4.2.

These results demonstrate that our algorithms improve sensitivity and specificity of the original anatomical parcellations. In the following, we will analyse features of our algorithms in more detail.

### 4.2 Effect of initial choice of parcellation

As can be seen in Fig. 5, some of the final parcels, particularly in the occipital, temporal and frontal lobes show overlaps across the three parcellations, while other regions in the central and parietal lobes can vary notably. All final parcellations in Fig. 5 are colour-matched to the first parcellation (modified Desikan-Killiany parcellation). To obtain a more direct comparison between the parcels, we computed the overlaps, normalised by the sizes of parcels (Fig. 6). More specifically, we took the modified Desikan-Killiany parcellation as the reference and found the overlaps between the colour-matched parcels in Fig. 5. Rows of the matrices in Fig. 6 illustrate the overlaps between each of the parcels of the parcellation on the y-axis (Py) with all the parcels of the parcellation on the x-axis (Px: always modified Desikan-Killiany), which is normalised by the size of that parcel of Py. Therefore, if there is only one yellow/white column corresponding to each row, it shows a one-to-one correspondence between the two intersecting parcels while several red/orange columns intersecting with each row show that one parcel in Py is overlapping with several regions in Px. If one row consists of only dark colours, that parcel in Py is not overlapping with any parcel in Px. As can be seen in Fig. 6, we found that a majority of parcels show a one-to-one correspondence between the final parcellations, with different degrees of overlaps. However, there are also several cases where a parcel in one parcellation overlaps with a few parcels or cases where a parcel does not have any matches in another parcellation.

**Figure 6.**
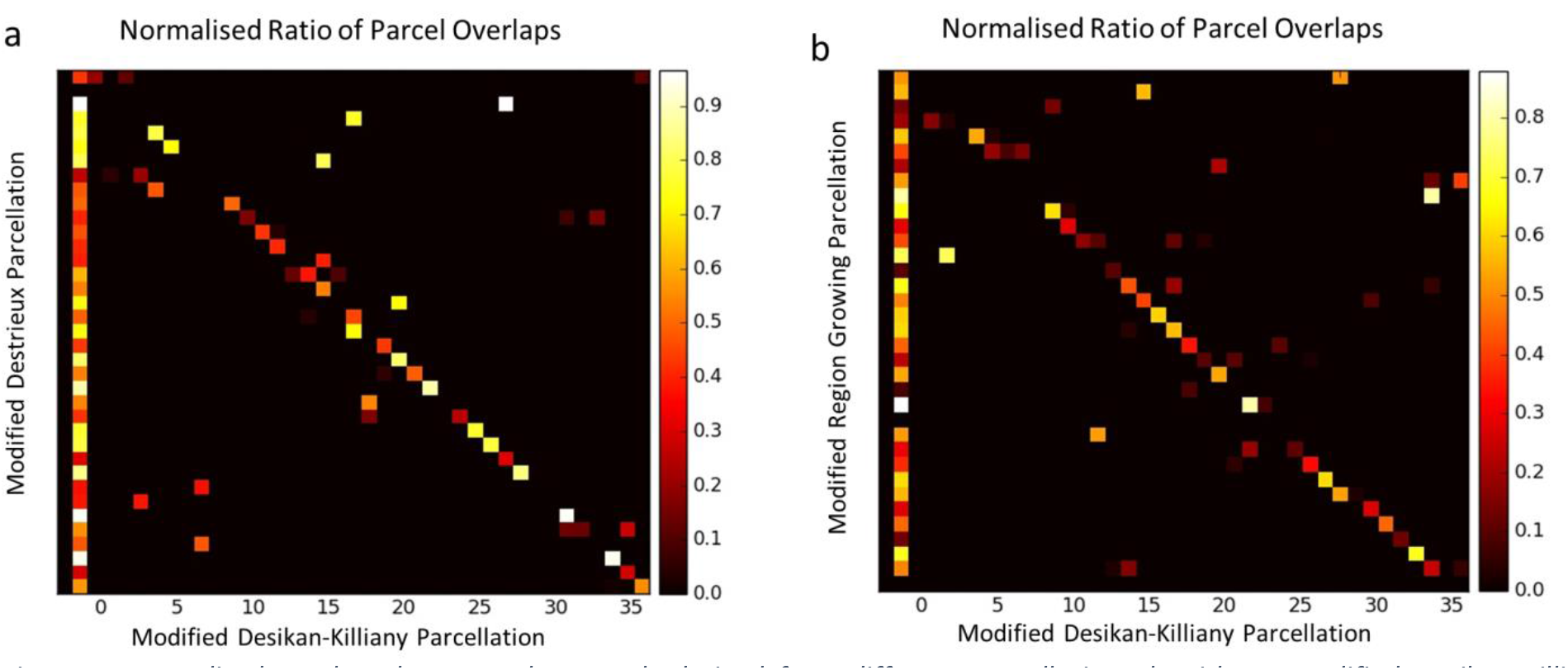
Normalised overlaps between the parcels derived from different parcellation algorithms. Modified Desikan-Killiany parcellation is shown on the x-axis and is used as the reference, (the order of parcels on the x-axis corresponds to Fig. 5a). Y-axis represents the parcels in a) modified Destrieux and b) RG parcellations. The rows correspond to the colour-matched regions of the x-axis and therefore the order is arbitrary in comparison to Fig. 5b, c. The sums of the normalised overlaps in each row are also shown as the first column.

### 4.3 Rank and condition number of final PRmats and implications

Here we compared the rank and condition numbers of PRmats for the original and modified parcellations. The resolution matrix, as expected, was highly ill-conditioned and while the ideal rank was 20484 in our study, the calculated rank was only 118. Parcellations (anatomical or modified) downsampled the source space to a few hundred parcels and thus improved the rank. We found a rank of 49 (ideal 68) and 92 (ideal 146) for the Desikan-Killiany and Destrieux atlases respectively, which, in spite of showing an improvement compared to the original source space, are still not full-ranked. In contrast, the modified parcellations showed near-perfect performance where we found ranks of 73 (ideal 74), 74 (ideal 74) and 70 (ideal 70) for the modified Desikan-Killiany, Destrieux and RG parcellations respectively. Even though full-ranked matrix guarantees independence between the parcel signals in the modified parcellations, the output might still be very sensitive to small changes in the input; hence a small condition number is desired. The condition numbers for the Desikan-Killiany and Destrieux atlases were 1.26x10^3^ and 1.78x10^4^ which were significantly improved to 114.38, 70.82 and 91.59 for the modified Desikan-Killiany, Destrieux and RG parcellations respectively. However, it is worth noting that condition numbers around 100 in the modified parcellations are still high and invite other complementary approaches to be used together with the EEG/MEG-adaptive parcellations. Some of these approaches will be discussed later.

### 4.4 Simulation results

In this section, we investigated the performance of anatomical and modified parcellations for realistic simulations of source networks where the ground truth is known, in order to address the following questions:

1. If there is no significant connectivity among the brain areas, how likely are different parcellations to identify significant false connections? These potential false connections will be merely leakage-induced and can act as a measure of susceptibility of each parcellation to leakage.
2. If networks with random active node (AN) locations and connections are simulated in the brain and different parcellations are used to reconstruct those networks, what is the accuracy of network reconstruction for each of the anatomical and modified parcellations?
3. Non-zero-lag connectivity measures such as imaginary part of coherency are insensitive to zero-lag connections. Leakage-induced connections are zero-lag. Does utilisation of non-zero-lag measures obviate the need for modified parcellations?

#### 4.4.1 Question 1: null networks

In order to compare different parcellations in the absence of true connectivity, we evaluated the null networks described in section ‎3.4.1.3. These networks include no active nodes and every vertex in the brain is given a random signal. Therefore, after network reconstruction and statistical analysis of the connectivity patterns, any significant connection is a false positive. It is worth noting that the False Positive Rate (FPR) of the simulated networks will have two underlying causes: 1-spurious connections due to leakage and 2-Type I error of statistical testing. The latter is corrected for multiple comparisons using permutation tests and is approximately 0.0001 (10000 permutations and uncorrected p-value of 0.05 (North et al. 2002)), for each simulated network, hence can be considered as a target FPR in the absence of leakage. Considering that this value is negligible compared to the observed FPR (Fig. 7), the main observed FPR for the null networks can be attributed to the leakage. Results are shown in Fig. 7. We computed the FPR for each null network by dividing the number of significant leakage-induced connections by the number of all possible connections for that network and computed the average FPR across 36 simulated datasets. The FPRs for Desikan-Killiany (DKA) and Destrieux (DA) atlases were 0.101 (231 out of 2278 possible connections corresponding to 68 nodes in DKA) and 0.081 (885 out of 10878 connections corresponding to 148 nodes in DA), respectively. The FPRs were reduced to 0.038 (101 out of 2701 possible connections for 74 parcels), 0.031 (85 out of 2701 connections) and 0.024 (60 out of 2415 connections for 70 parcels) for modified DKA, modified DA and RG respectively. Therefore, the modified parcellations’ FPRs were about one third of those of the anatomical parcellations.

**Figure 7:**
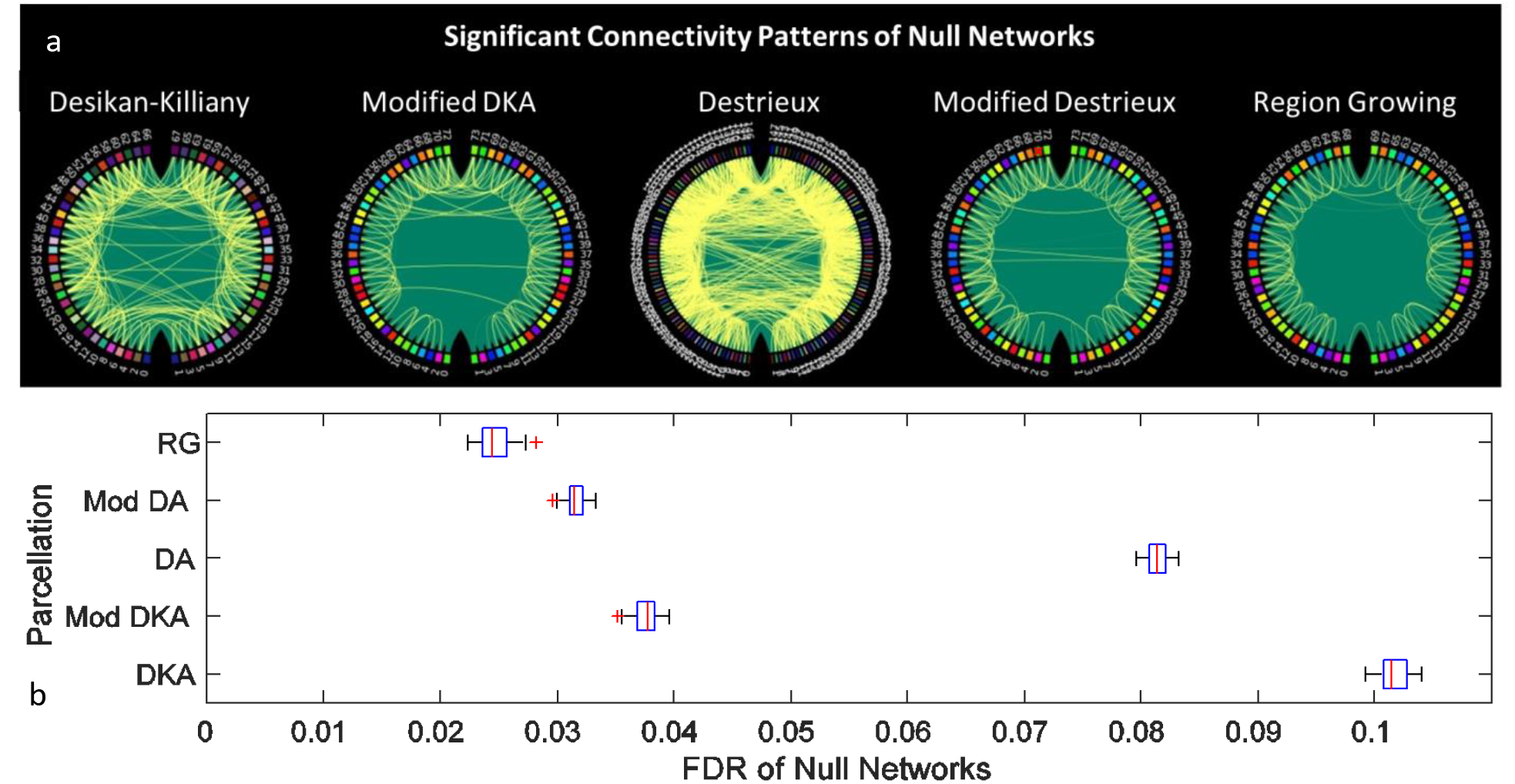
a) Significant connections of the null networks for the anatomical and modified parcellations. The ratio of the leakage-induced connections (false positives) to all possible connections was found to be 0.101 and 0.081 in DKA and DA atlases that was decreased to 0.038, 0.031 and 0.024 in the modified DKA, modified DA and RG parcellations respectively. Node colours correspond to the node colours in Figs. 3, 4. b) Variations of FDRs for null networks of each parcellation across 36 simulated datasets (boxplots mark median (red lines), standard deviations (in blue), confidence interval (in black) and outliers (red cross).

#### 4.4.2 Question 2: realistic networks with active nodes

We simulated hundreds of realistic datasets with varying numbers of ANs (3, 5, 10, 15), percentage of connections among ANs (25%, 50% and 100%) and SNR of the data (1.0 and 3.0). For each of these scenarios, 36 datasets each consisting of 17 subjects were simulated where the locations of ANs and connections randomly varied across datasets. ANs were random parcels selected from Brainnetome functional atlas (Fan et al. 2016, Fig. B. 1, Appendix B). Thereafter, we used bivariate coherence for network reconstruction and identified significant connections of each network across subjects using permutation tests. Significant connections of one example network with 5 ANs and 100% connections among the ANs is shown in Figure 8. It is worth noting, since the ANs are based on the Brainnetome atlas, each AN might show spatial overlap with several nodes in each parcellation and therefore the number of active parcels found in any simulated network might be higher than the number of ANs. Furthermore, some of the ANs did not overlap with any of the modified parcels due to the lower coverage of the cortex by these adaptive parcellations (Fig. B. 1). Table 2 presents the average number of missed ANs and average number of parcels per AN for each parcellation. Approximately 20-30% of connections were missed due to no coverage by modified parcellations and average parcel per AN for anatomical parcellations was approximately two times that of modified parcellations.

**Figure 8.**
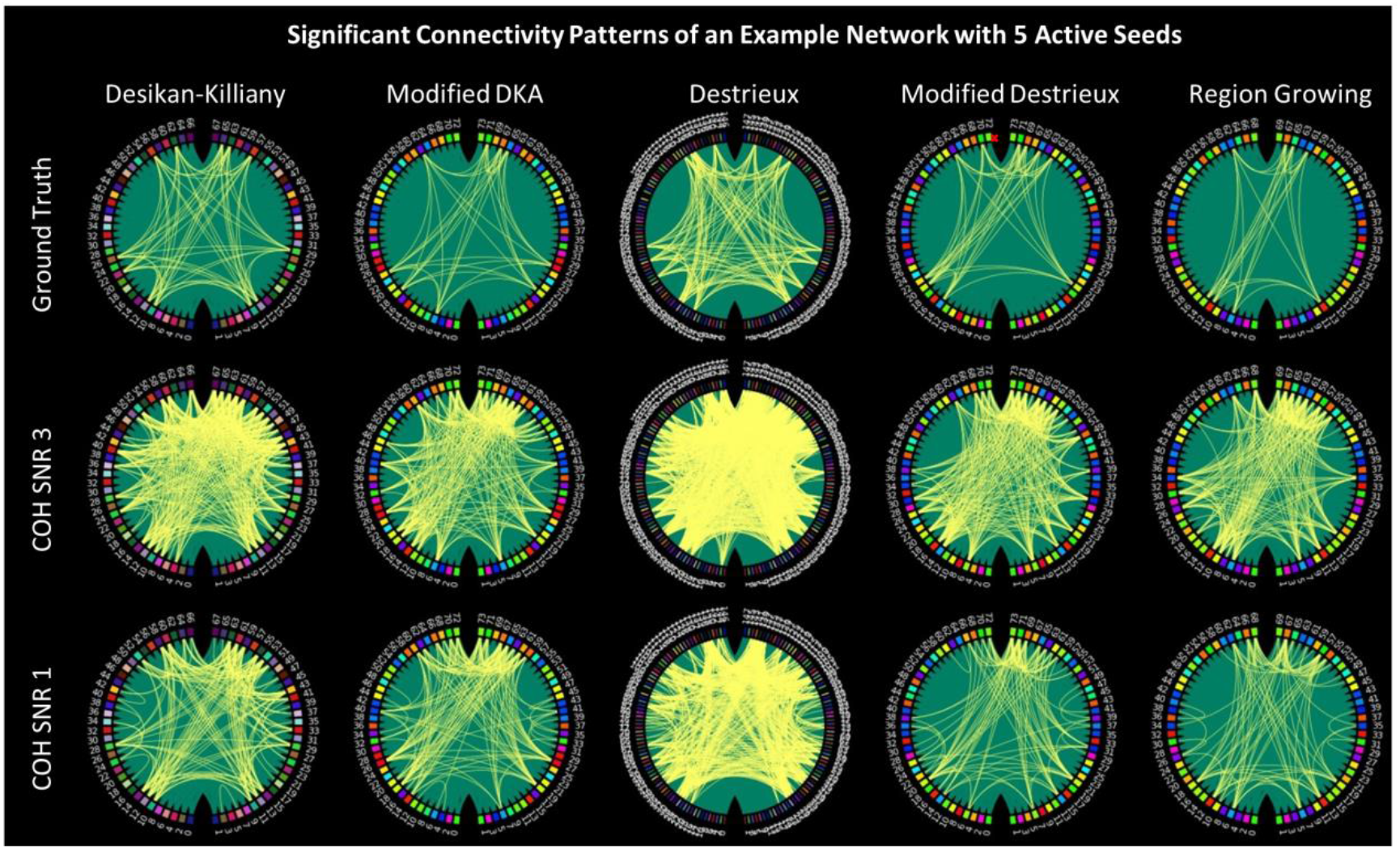
Significant connections for an example network with 5 active seeds. The first row shows the ground truth in the absence of leakage, the second and third rows show the network in the presence of leakage under SNR 3 and 1 respectively, as computed using coherence. Node colours correspond to the node colours in Figs. 3, 4.

**Table 2.** Average ratio of missed nodes (no overlap between an AN from Brainnetome atlas and parcels of a parcellation), missed connections due to the missed nodes and parcels per AN across all the simulated scenarios with different number of ANs and connections. DKA: Desikan-Killiany Atlas, DA: Destrieux Atlas, RG: Region Growing, AN: Active Node.

|  | DKA | Mod DKA | DA | Mod DA | RG |
| --- | --- | --- | --- | --- | --- |
| **Missed Nodes** | 0 | 0.19 | 0 | 0.26 | 0.28 |
| **Missed Connections** | 0 | 0.19 | 0 | 0.26 | 0.30 |
| **Parcels per AN** | 2.91 | 1.63 | 3.87 | 1.4 | 1.41 |

Here, we compared parcellation-specific ground truths (e.g. first row in Figure 8) in the absence of leakage to the realistic networks in the presence of leakage. Fig. 9 summarises the proportions of true positive (TPR) and false positive rates (FPR) for each of the parcellations and in each of the simulated scenarios. Note that since the modified parcellations in Fig. 5 do not cover some of the functional seeds of the Brainnetome Atlas (Fig. B. 1) and the locations of active seeds are selected randomly, some seeds are missing in both ground truth and realistic scenarios of the adaptive parcellations. We identified the missing seeds and the corresponding connections for each modified parcellations using the procedure described in 3.4.2.1 and included them in the computation of TPRs. Thus, a parcellation that covers the

**Figure 9:**
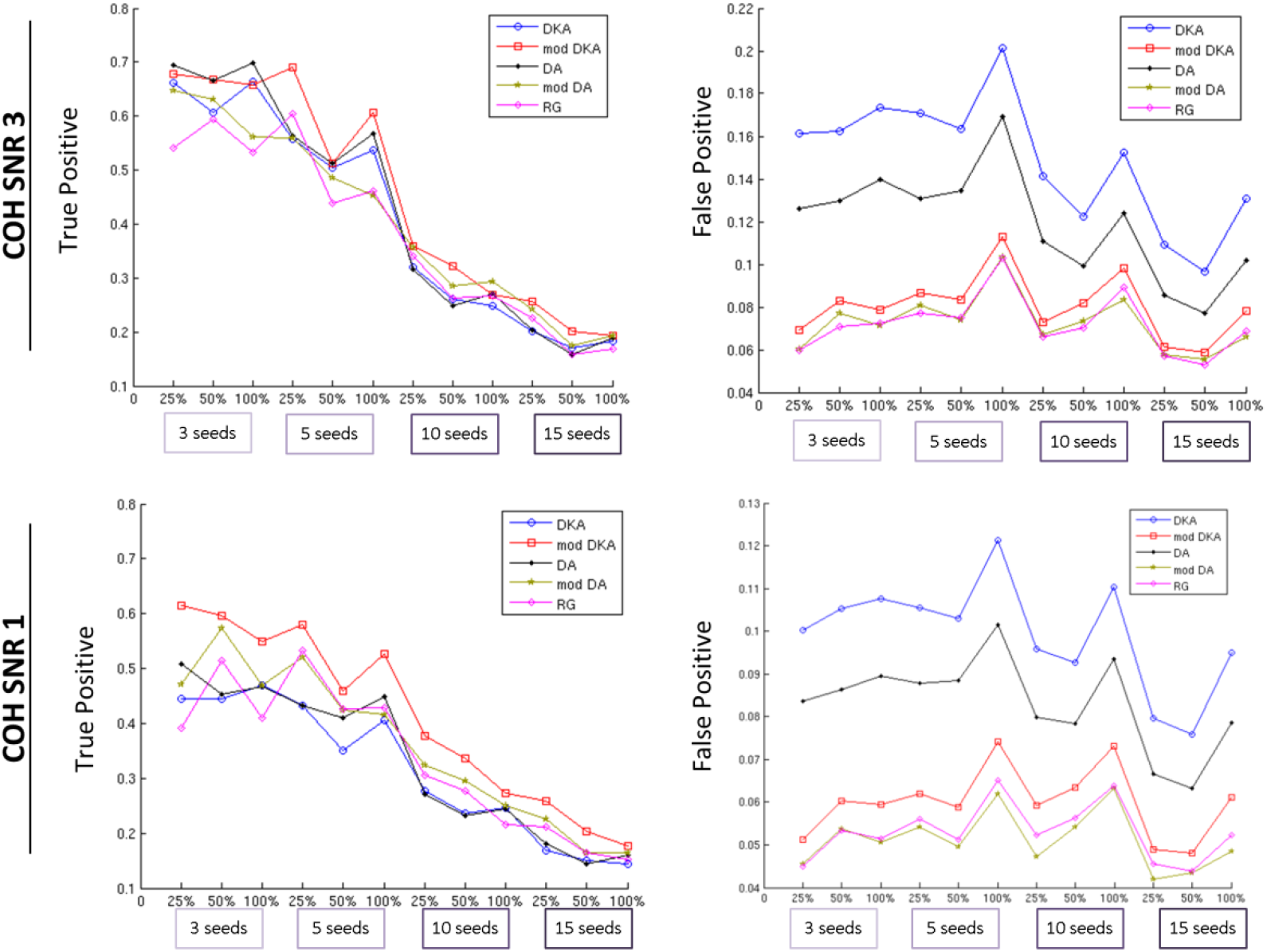
True positive and false positive rates obtained from coherence analysis of networks based on the anatomical and modified parcellations at SNRs 1.0 and 3.0. The reference for each parcellation is the reconstructed network in the absence of leakage. TPRs and FPRs are obtained by comparing each reconstructed network in the presence of leakage to the reference network for the same parcellation. COH: Coherence, SNR: Signal-to-noise ratio, DKA: Desikan-Killiany atlas, DA: Destrieux atlas, mod: modified, RG: Region Growing.

AN in the Brainnetome atlas may have a chance to recover it, which would contribute to its TPR. However, if it mislocalises activity from this AN or is insensitive to it, it will count towards the FPR or reduce TPRs. For a parcellation that does not cover the AN at all, it will automatically contribute to (reduction of) TPR. Table 3 presents the details of statistical comparisons between average TPRs and FPRs of each pair of parcellations across different numbers of seeds and connections. The p-values in this table are obtained using Kruskalwallis test which is the non-parametric ANOVA in order to account for the potential non-normalities in the distributions of the variables.

**TPR**: As shown in Fig. 9, at SNR 3, TPRs were reduced as the number of seeds/connections increased. For the anatomical parcellations, starting from ~0.7 true positives for 3 seeds, TPR was reduced to ~0.5 for 5 seeds, and then dropped sharply to 0.3 or less for 10 and 15 seeds. We found a similar trend for the modified parcellations, except that on average, modified DKA showed significantly higher TPRs than anatomical parcellations (c.f. Table 3 for details). We found no significant improvements for the TPRs of modified DA and RG compare to anatomical parcellations. At SNR 1, we found a similar trend, but not surprisingly (as elaborated in Table 3), the TPRs at SNR 1 where on average lower than SNR 3, for both anatomical and modified parcellations. Furthermore, as shown in Fig. 9, the gap between TPRs of anatomical and modified was larger for SNR 1 compared to SNR 3. Additionally, both modified DKA and DA showed significantly higher TPR compared to anatomical parcellations.

**FPRs:** Even though TPRs were improved for some of the modified parcellations, we observed the main improvements in the FPRs. Anatomical parcellations showed substantially lower specificity compared to all the modified parcellations, by a factor of 2 or more (Table 3). Within anatomical parcellations, DKA showed significantly higher FPRs compared to DA. Furthermore, modified DKA showed significantly higher FPR compared to modified DA and RG. Moreover, we found that for 5, 10 and 15 seeds, FPR peaked when 100% of connections among seeds were present, suggesting that fuller networks might suffer more from leakage-induced false positives. At SNR 1, we found very similar trends as SNR 3 for both anatomical and modified parcellations. FPR was reduced at SNR 1, probably because low SNR results in larger variability and generally decreased detection of connections.

Therefore, we observed a substantial improvement in the FPRs and some improvement in the TPRs. It is worth noting that since modified parcellations do not cover some of the functional seeds of the Brainnetome Atlas (Fig. B. 1), two separate factors influence TPRs: 1-missing seeds; 2-sensitivity score for the areas that are covered by the modified parcellations. In order to tease these two factors apart and focus on number 2, we also computed TPRs without taking the missing sources into account. This was done by comparing each network in the presence and absence of leakage (e.g. first vs. second rows of Fig. 8). Results are shown in Fig. B. 2 and Table B. 1 in Appendix B. These results are informative since they show the sensitivity in the areas covered by each parcellation (Fig. B. 1) and additionally correspond to the parcellation performance indices provided by the PRmats that are also based on the “covered parcels” only.

**Table 3.** Statistical comparison of average TPRs and FPRs of anatomical and modified parcellations. Blue shading: adaptive significantly improved compared to anatomical. Orange shading: Higher performance in anatomical. Grey shading: Significant differences among the adaptive parcellations. Green shading: best average performance.

| True Positives |  |  |  |  |  |  | False Positives |  |  |  |  |  |  |
| --- | --- | --- | --- | --- | --- | --- | --- | --- | --- | --- | --- | --- | --- |
| Coherence SNR 3 |  |  |  |  |  |  | Coherence SNR 3 |  |  |  |  |  |  |
|  | mean±std | p-value |  |  |  |  |  | mean±std | p-value |  |  |  |  |
|  |  | DKA | DA | Mod DKA | Mod DA | RG |  |  | DKA | DA | Mod DKA | Mod DA | RG |
| DKA | 0.409±0.055 | NA | 0.21 | 0.002 | 0.87 | 0.07 |  | 0.149±0.011 | NA | 1.65E-12 | 2.92E-13 | 2.92E-13 | 2.92E-13 |
| DA | 0.424±0.044 | 0.21 | NA | 0.027 | 0.40 | 0.002 |  | 0.119±0.01 | 1.65E-12 | NA | 2.92E-13 | 2.92E-13 | 2.92E-13 |
| Mod DKA | 0.451±0.054 | 0.002 | 0.027 | NA | 0.006 | 1.24E-05 |  | 0.08±0.006 | 2.92E-13 | 2.92E-13 | NA | 6.30E-06 | 9.63E-07 |
| Mod DA | 0.407±0.065 | 0.87 | 0.40 | 0.006 | NA | 0.07 |  | 0.073±0.007 | 2.92E-13 | 2.92E-13 | 6.30E-06 | NA | 0.67 |
| RG | 0.383±0.061 | 0.07 | 0.002 | 1.24E-05 | 0.07 | NA |  | 0.072±0.006 | 2.92E-13 | 2.92E-13 | 9.63E-07 | 0.67 | NA |
| Coherence SNR 1 |  |  |  |  |  |  | Coherence SNR 1 |  |  |  |  |  |  |
| DKA | 0.314±0.047 | NA | 0.26 | 1.37E-09 | 0.001 | 0.21 |  | 0.099±0.008 | NA | 4.09E-10 | 2.92E-13 | 2.92E-13 | 2.92E-13 |
| DA | 0.33±0.044 | 0.26 | NA | 2.64E-08 | 0.015 | 0.87 |  | 0.083±0.008 | 4.09E-10 | NA | 4.08E-13 | 2.92E-13 | 2.92E-13 |
| Mod DKA | 0.413±0.054 | 1.37E-09 | 2.64E-08 | NA | 0.002 | 7.00E-06 |  | 0.06±0.007 | 2.92E-13 | 4.08E-13 | NA | 2.01E-06 | 8.48E-05 |
| Mod DA | 0.359±0.073 | 0.001 | 0.015 | 0.002 | NA | 0.11 |  | 0.051±0.006 | 2.92E-13 | 2.92E-13 | 2.01E-06 | NA | 0.23 |
| RG | 0.336±0.065 | 0.21 | 0.87 | 7.00E-06 | 0.11 | NA |  | 0.053±0.006 | 2.92E-13 | 2.92E-13 | 8.48E-05 | 0.23 | NA |

#### 4.4.3 Question 3: imaginary part of Coherency

In this section we investigated whether or not non-zero-lag connectivity measures such as imaginary part of coherency will also benefit from E/MEG-informed parcellation modifications. Fig. 10 displays the TPR and FPRs for the ImCOH (similar to Fig. 9 for Coherence). We found TPR and FPR patterns that resembled the Coherence results. FPRs were generally higher for the anatomical compared to the modified parcellations. TPRs did not differ between the anatomical and modified parcellations. Therefore, modified parcellations also improve the ImCOH results. Both TPRs and FPRs were lower for ImCOH compared to COH. Lower TPR is presumably due to the fact that true zero-lag connections were not detectable using ImCOH, and ImCOH also attenuates near-zero-lag connections. Additionally, lower TPRs and FPRs might be attributed to the fact that ImCOH ignores the real part of the signal and thus yields smaller connectivity values that are more likely to become obscured by the presence of noise in the data.

**Figure 10.**
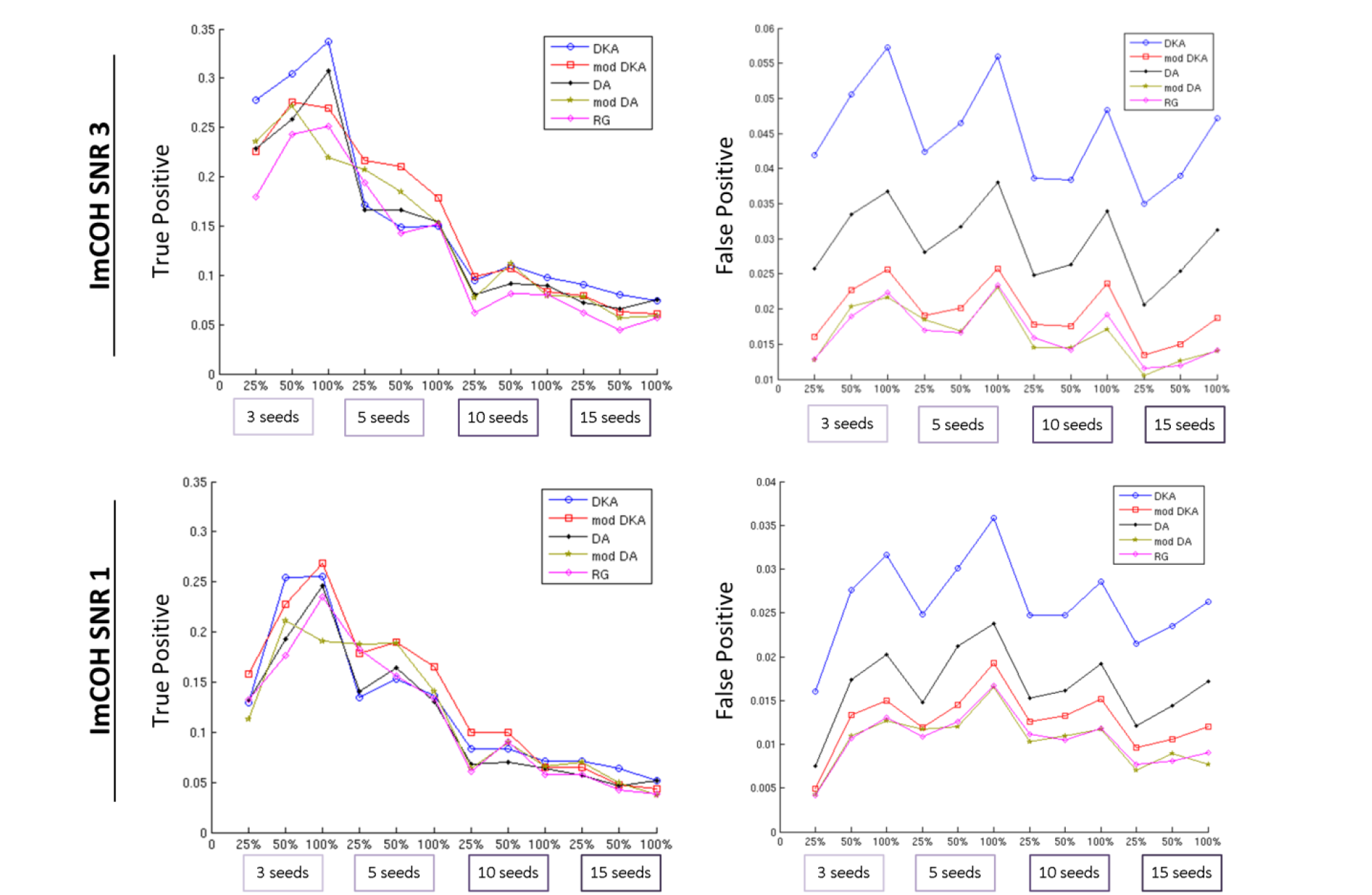
True positive and false positive rates obtained from ImCOH analysis of networks based on the anatomical and modified parcellations at SNRs 1.0 and 3.0

## 5 Discussion

We used cross-talk functions (CTFs), which describe the spatial resolution of linear or linearly constrained distributed source models, to create EEG/MEG-adaptive parcellations of the cortex as a basis for connectivity analysis of EEG/MEG data in source space. We implemented two CTF-based algorithms– split-and-merge (SaM) and region growing (RG) – which differed with respect to the starting points of the parcellation process. For SaM, we started from two different standard anatomical parcellations with different average sizes of parcels (Desikan-Killiany (Desikan et al. 2006) and Destrieux (Destrieux et al. 2010) Atlases) and modified the parcels so as to comply with the spatial resolution of EEG/MEG. For RG, we started with no prior parcellation and created a parcellation from all the brain vertices. We used metrics for distinguishability and sensitivity based on parcel resolution matrices (PRmats) to quantify the performance of different parcellations, using a dataset consisting of combined EEG and MEG measurements. All three analyses yielded approximately 70 distinguishable parcels in the brain, suggesting that this reflects the general resolution limits of the utilised measurement configuration and source estimation methods. All approaches provided a sparse sampling of the cortex, and significantly improved the parcellation performance compared to the anatomical parcellations with respect to sensitivity and distinguishability of parcels, while at the same time maximising the number of distinguishable parcels in the brain.

Furthermore, using extensive realistic simulations, we showed that: a) when there are no true connections among the brain areas, the ratio of false leakage-induced connections detected by the modified parcellations were improved by a factor of two or more compared to the anatomical parcellations; b) in the presence of active sources and connections, adaptive parcellations showed higher sensitivity and substantially higher specificity (up to improvement factor of 2); and c) modified parcellations improved the network reconstruction accuracy using both zero-lag and non-zero-lag connectivity measures.

### 5.1 Adaptive parcellations for the spatial limitations of EEG/MEG

EEG/MEG studies typically adopt anatomical or fMRI-based functional parcellations. For example, Hillebrand et al. (2012) used the Talairach Daemon Database for the parcellation of the brain, Colclough et al. (2015, 2016) used the Harvard-Oxford anatomical parcellation and ICA-based fMRI parcellation, while several other studies have used the Automatic Anatomical Labelling (AAL) atlas (Tewarie et al. 2014; Tewarie et al. 2016; Brookes et al. 2016). Nevertheless, as described in the theory section, anatomical parcels are unlikely to be optimal for EEG/MEG analysis. Therefore, in the current study we introduced EEG/MEG-adaptive parcellations. For this purpose, we used a state-of-the-art measurement configuration containing EEG and MEG sensors, realistic individual boundary element (BEM) models (Fuchs et al. 2002) and L2 minimum norm (MNE) source estimation that makes minimal assumptions about the source configuration (Hämäläinen & Ilmoniemi. 1994; Hauk 2004) and introduced novel approaches for parcellating the cortex.

The parcellation algorithms implemented here are adaptive and can change depending on the choices of EEG/MEG measurement configuration, head model and source estimation methods. Therefore, since it has been shown previously that combining EEG and MEG provides higher spatial resolution (Fuchs et al. 1998; Molins et al. 2008; Henson et al. 2009) it can be expected that EEG or MEG on their own will result in a smaller number of surviving parcels than for their combination. Moreover, we have used a common BEM model in our forward computations (Hämäläinen & Sarvas 1989; Mosher et al. 1999). It can be expected that using other multi-layer headmodels or Finite Element Models (FEMs) (Buchner et al. 1997) may also change the parcellations. Furthermore, different source estimation methods will result in different CTFs. It is important to note that due to Equation 5, all CTFs, regardless of the inverse methods used, are linear combinations of the leadfields. Thus, CTFs that are not in the space of the leadfields cannot be achieved by any method. In this study, we used L2 MNE that results from the minimisation of the difference between the resolution matrix and the identity matrix (Dale & Sereno 1993; Hauk 2004) and yields an optimum source localisation when no further specific modelling constraints are applicable. However, it is worth noting that a bias towards superficial sources is often associated with L2 MNE which might have resulted in the exclusion of deeper brain areas in the adaptive parcellations. This bias is present in the leadfields of EEG and MEG sensors, and will therefore be inherited by CTFs for all linear methods. Nevertheless, in studies where other constraints are justified, e.g. when other families of spatial filters such as beamformers (Van Veen et al. 1997; Barnes et al. 2006) or weighted MNE are used, different parcellations of the cortex and possibly more coverage of deeper brain areas can be expected. It is also worth noting that in the current study with unweighted L2 MNE source localisation which depends on the data only through regularisation by the noise covariance matrix, the final number of parcels in a parcellation is relatively independent of the degrees of freedom in the data. However, the final number of parcels would be expected to have stronger data-dependence if e.g. beamformers were used.

#### 5.1.1 Comparison to the precedent EEG/MEG-informed parcellations

A few previous studies have also investigated EEG/MEG-informed parcellations. To the best of our knowledge, Palva et al. (2010) presented the first study that has used an EEG/MEG-informed parcellation. They utilised the forward and inverse modelling of simulated noise in source space and by means of k-means clustering identified 365 patches on the cortex that showed high within-patch phase synchrony. This method has several advantages including a) yielding parcellations based on EEG/MEG data; b) yielding individualised parcellations that are suitable for single subject connectivity analysis and c) unlike CTF-based approach used in the current study, it is not restricted to the linear/linearly constrained distributed source models. However, there are a few disadvantages that we have tried to overcome in the current study: 1) in Palva et al.'s study, the parcellation was done at single-subject level and therefore the locations of different parcels can vary significantly across the subjects. This makes the method unsuitable for group analysis; 2) the number of parcels in the brain were fixed in that study at the same number as the utilised sensors. This is arguably an arbitrary choice for the number of spatially independent parcels and thus might yield some highly dependent parcels on the cortex (the parcel sensitivity and specificity was not reported). However, the spatial resolution of EEG/MEG is confined by the measurement configurations, head models and inverse operators. CTFs quantify these spatial limitations and therefore can be used to determine the number, location and sizes of independent parcels; 3) in that study, the parcels were determined based on forward and inverse modelling of random noise. Therefore, the identified patches can be good representatives of independent patches for the real datasets only if the simulated noise properties provide a good model of the real data; 4) the parcels were determined based on a specific connectivity metric (e.g. phase-locking values). The choice of connectivity metric can change and the algorithms will adapt to different choice of connectivity metrics and yield connectivity-dependent parcellations. However, if a comparison of the results of network reconstructions using several connectivity metrics is of interest in one study or comparison between different studies, Palva et al.'s algorithm will yield different parcellations for different metrics which makes this comparison difficult.

In a more recent study, Korhonen et al. (2014) introduced sparse weights to collapse the source space, based on the forward and inverse modelling of simulated noise in the source space, so that vertex selection is optimised for a fixed set of predefined anatomical parcels. Their method, likewise Palva et al.'s, has the advantage of being generalisable to the source models that are not distributed and linear/ linearly constrained. Furthermore, their method is suitable for group-level analysis while it takes the individual differences into account. This is achieved by fixing the number and locations of parcels based on the anatomical atlases while selecting representative vertices for each parcel based on the forward and inverse models of each individual. However, the disadvantages 2-4 mentioned above for Palva et al.'s study are also applicable to the sparse weights approach. Therefore, obtaining a parcellation that can overcome these problems and at the same time optimises both parcellation resolution (i.e. number of parcels in a parcellation) and vertex selection with respect to EEG/MEG spatial limitations, had remained a challenge (Korhonen et. al 2014) which we have tried to overcome in the current study.

### 5.2 Different parcellation approaches: similarities and differences

Our proposed parcellation algorithms addressed the three theoretical issues of using anatomical parcels with EEG/MEG that were discussed in the Theory section (Fig. 1). Firstly, adaptive parcellations identified and ignored vertices that our source estimation methods provided a low sensitivity to. More specifically, we found a limited sensitivity to the signals that are produced in deeper brain areas. All three parcellations (Fig. 5) showed almost no coverage of the medial view of the cortex indicating the relative insensitivity of our source estimation to these deeper brain cortices. It is worth noting that while the locations of the cortices with low sensitivity might change depending on measurement configuration and source localisation, our results suggest that the proposed algorithms can identify those vertices successfully. Secondly, the specificity of the anatomical parcellations did not match that of the EEG/MEG parcellations: On the one hand, some fine-grained neighbouring areas were not distinguishable. For example, the four areas pars-triangularis, pars-orbitalis, pars-opercularis and lateral orbitofrontal cortex from the Desikan-Killiany atlas (Fig. 4a) were merged into two areas in the anterior and posterior inferior frontal gyrus in the modified version of this atlas (Fig. 5a). On the other hand, large parcels such as pre- and post-central gyri were split into smaller parcels (e.g. compare Fig. 4a and Fig. C. 1a).

The two SaM and RG approaches showed highly overlapping final parcels for all three final parcellations, which indicates the robustness of the proposed algorithms with respect to the initial choice of parcellation. This indicates that the final parcellation of the cortex is mostly influenced by the choices of measurement configuration, head model and source estimation method. However, as shown in section 4.2, we observed notable differences as well, in that not all the parcellations provide a similar sparse sampling of all the brain areas. For example, as can be seen in Fig. 5, while the final RG parcellation includes several parcels in the temporal lobe, the modified Destrieux parcellation provides a better coverage of centro-parietal cortices. Furthermore, as shown using simulated networks and as a result of different samplings of the cortex, modified parcellations provided different sensitivity and specificity of network reconstructions. More generally, the SaM approach is based on anatomically defined regions and thus provides a better solution for optimising the number of a priori selected parcels or testing specific hypotheses. In contrast, the RG approach is most distinct from anatomical labels and limitations that they could impose on detection of functional networks. Therefore, it might be more desirable for data-driven whole brain connectivity analyses, e.g. for resting state networks.

### 5.3 Effect of parcellations on reconstruction of realistic networks

Our simulations were set up to investigate the effects of different parcellations on the accuracy of network reconstructions in several scenarios by varying the number and locations of active sources in the brain, percentage of connections among those sources as well as SNR of the data. Active sources were defined by randomly selecting functional parcels from the Brainnetome atlas (Fan et al. 2016) so as to obtain a realistic representation of the size and locations of functional nodes in the brain. We found that adaptive parcellations show up to three times less false leakage-induced connections among the parcels. This was found by investigation of a) null networks with realistic levels of noise and no active sources as well as b) realistic networks with multiple active sources. Furthermore, we observed improvements in detection of true connectivity among the sources. This was interestingly in spite of the fact that approximately 20% of connections were missed by the modified parcellations since the corresponding seeds of the Brainnetome atlas were not covered by the modified parcellations and thus the maximum true positive rates that could have been achieved using adaptive parcellations were approximately 0.8. Even so, the overall true positives detected by the modified parcellations were at a same level or higher than the anatomical parcellations. This was particularly evident for lower SNR of the data (SNR 1.0 compared to 3.0), suggesting that optimal parcellation is more crucial in the presence of higher levels of noise. Therefore, our investigation of sensitivity and specificity depict that if the spatial resolution of EEG/MEG and source localisation methods do not allow for inclusion of some the brain cortices, including them in the model will reduce the specificity substantially while it does not allow for improvements in sensitivity.

Comparing the performance of different adaptive parcellations, we found that modified DKA showed higher sensitivity compared to modified DA and RG while modified DA and RG showed higher specificity. Modified DKA is based on the Desikan-Killiany atlas which includes 68 parcels that is of the same order as the final adaptive parcellations. In other words, it can be considered a good starting point for initiation of the parcellation algorithms. Therefore, the modification procedure resulted in fewer changes in this parcellation and higher overall coverage compared to modified DA and RG. In contrast, modified DA and RG yielded the most distinct parcels in the brain and higher specificity at the expense of less coverage. Therefore, even though as discussed in 5.2 different parcellations provided highly overlapping results, differences in network reconstructions suggest that depending on the purpose of a study and locations of the networks of interest, one initial point and/or algorithm might prove more useful.

Another finding involved the trend of changes in the sensitivity and specificity as a function of changes in the number of simulated active sources/connections in the brain. The sensitivity for all anatomical and adaptive parcellations decreased with increases in the number of active sources/connections in the brain. In particular, we observed a sharp change from 5 seeds to 10 seeds. This shows that the accuracy of network detection drops substantially for widespread or highly dense networks. However, it is worth mentioning that the number of seeds is not equal to the number of active parcels since the seeds are derived from the Brainnetome atlas and parcels of this atlas might overlap with several parcels in the parcellations. As a matter of fact, 10 active seeds might actually correspond to 20 active parcels or more. On the other hand, the false positives showed a trend that was similar among different parcellations. Unlike true positive rates, FPRs were fluctuating more depending on the percentage of connections among the seeds rather than the number of active nodes. For example, keeping 100% of connections resulted in higher FPRs compared to 25% of connections for any number of seeds, suggesting that fuller networks are more prone to the leakage problem.

### 5.4 Non-zero-lag connectivity does not obviate the need for EEG/MEG-adaptive parcellation

Non-zero-lag connectivity measures have been introduced to alleviate the leakage problem (Nolte et al. 2004; Stam et al. 2007). We investigated whether using non-zero-lag connectivity can resolve the need for an adaptive parcellation for whole-brain network analysis. We used magnitude-squared coherence (COH) and imaginary part of coherence (imCOH) as spectral measures of synchrony (Greenblatt et al. 2012; Bastos & Schoffelen 2016). While COH is sensitive to zero- as well as non-zero-lag connections, imCOH is only sensitive to the latter. We argued (section 2.1.4) that even bivariate and multivariate non-zero-lag connectivity measures are affected by leakage. Furthermore, we showed that long-range spurious connections between a seed and a target can occur due to leakage to the target (i.e. inherited connectivity (Colclough et al. 2015)). By means of realistic simulations we found that ImCOH: a) resulted in fewer false positives as expected but did not resolve the FPR problem. In fact, FPRs obtained from imCOH for anatomical parcellations were comparable to FPRs obtained from COH using adaptive parcellations; b) showed substantially lower FPRs for modified compared to anatomical parcellations and c) showed notably less TPRs for both anatomical and modified parcellations. The latter can be attributed to the fact that imCOH does not detect true zero-lag connections and attenuates nearzero-lag connections. Additionally, lower TPRs and FPRs might be attributed to the fact that ImCOH ignores the real part of the signal and thus yields smaller connectivity values that are more likely to become obscured by the presence of noise in the data. Therefore, it appears that using imCOH with anatomical parcellations can reduce false positives at the expense of lowering true positives while utilising COH with adaptive parcellations can result in comparable reductions in FPRs without compromising the TPRs. Additionally, if having low FPRs is of main interest in a study, or zero-lag connections are assumed unlikely or irrelevant in a dataset, combining imCOH with adaptive parcellations can result in a high suppression of FPRs.

### 5.5 Practical notes

Here we discuss two practical considerations. Firstly, the parcellation introduced in this study is defined in a standard source space where CTFs computed in the single subject space are morphed to a standard space and averaged across a group of subjects for further analysis. These standard parcels can be morphed to the individual source spaces for the single subject analysis. In a series of trials that are not reported here, we found that parcels defined in single subject space can be highly inconsistent across subjects. This is firstly due to the fact that the sizes of parcels can vary largely across subjects and secondly, some overlapping vertices might be assigned to different anatomical labels in different subjects. Therefore, we conclude that in order to obtain a consistent set across subjects and robustness to noise, parcels can be defined in a standard canonical space and, if single subject connectivity analysis is of interest, parcels can be morphed to the individual source space.

Secondly, there are several SaM and RG parameters that can be adjusted in order to obtain a parcellation that is most suitable for the questions of a study; here we used generalisable parameters based on the values commonly used for similar purposes in the literature. First, in order to assign a vertex to a parcel we used half-maximum of CTF values as a measure of sensitivity. Half-maximum is commonly used in signal processing as a measure of sensitivity in order to provide a cut-off to assign a set of values to a given peak. In signal processing, it corresponds to ~3dB attenuation in the power of the signal (below which the signal is considered damped) (Oppenheim & Schafer. 2010). Second, we used z-scores above 3 for sensitivity and specificity of a vertex to a parcel. A Z-score above 3 for Gaussian distributions corresponds to a p-value<0.005, showing that a vertex is significantly more sensitive to a given parcel as compared to any other parcels. Third, we allowed at least one standard deviation between the parcel with highest specificity and the parcel with next highest specificity (see Methods section for details). These values can be considered “standard” to provide a reasonable tradeoff between the sensitivity, specificity and maximising the number of distinguishable parcels. However, if there are clear requirements for sensitivity versus specificity, these values can be adjusted to adapt the parcellation accordingly. Another parameter is the minimum number of vertices that are required to form a separate parcel. We heuristically selected a minimum of 10 vertices, in order to exclude very small parcels that might be significantly affected by slightly changing other parameters of parcellation.

### 5.6 Future directions

In this study we showed that obtaining EEG/MEG-adaptive cortical parcellations can improve sensitivity and distinguishability of the parcels in the brain, both in theory using resolution matrices and in practice using simulated networks. However, it is worth noting that the proposed algorithms depend on the choice of forward model and inverse solution and are consequently sensitive to the potential modelling errors. The degree to which the proposed algorithms are sensitive to the modelling errors can be an interesting scope for the future research. Moreover, the current implementations of our algorithms are suitable for studies on homogeneous cohorts of participants. This is due to the fact that depending on the choice of source localisation methods, adaptive parcellations might be more or less data-dependent. For example, L2 MNE depends on the noise-covariance of the data and is somewhat data-dependent (depending on the regularisation parameter) and beamformers depend on the data covariance and are strongly data-dependent. Therefore, if the relevant data properties (e.g. noise covariance or data covariance) change significantly between different groups under investigation (e.g. patients and healthy participants), different parcellations might be obtained for different groups. This can result in significant differences in the functional networks. Thus, generalisation of current approaches to inhomogeneous groups could be explored in the future studies. These outstanding questions provide exciting opportunities for the future research on adaptive parcellations.

Furthermore, the final parcel resolution matrices and simulation results suggest that network reconstruction accuracy has been notably improved in adaptive parcellations. However, the presence of off-diagonal elements in PRmats (specifically reflected in the results of simulated null networks) allow for using adaptive parcellations together with complementary methods from the previous literature that can be expected to further improve network reconstruction accuracies. One such complementary method might be to combine adaptive parcellations with multivariate connectivity. In the theory section and Appendix A, we have discussed how multivariate and non-zero-lag connectivity methods can be affected by the leakage, and considering the linear nature of CTFs and based on the multivariate covariance as an example, we discussed that leakage coefficients could be taken into account in order to quantify the effects of CTFs on multivariate connectivity analysis. These leakage coefficients can be extracted from the PRmats. Therefore, we suggest that modified parcellations and PRmats might be used together with multivariate and time/phase-lagged estimates of connectivity, in order to get more direct and directed measures of whole-brain graphs. It is worth noting that computing PRmats for any given parcellation (e.g. anatomical parcellations) to inform the multivariate connectivity analysis might not result in an accurate reconstruction of whole-brain networks. This is due to the fact that standard anatomical parcellations are likely rank-deficient (section 4.3) which indicates that signals of one or more parcels are strongly dependent on a linear combination of other parcels in the brain and cannot be estimated accurately. On the contrary, the parcellation algorithms in this study improved this issue substantially, suggesting that one can derive N independent signals for N parcels yielded by the parcellation algorithms. Therefore, obtaining distinguishable CTF-based parcels is an essential first step and how to combine these adaptive parcellation methods with different connectivity methods will be an important question for future studies.

## 6 Acknowledgements

This work was supported by a Cambridge University international scholarship award to S.F and UK Medical Research Council grants to R.N.H. (MC_A060_5PR10) and O.H. (MC_A060_53144). We would like to thank Dr. Darren Price for commenting on an earlier version of this manuscript and Dr. Karalyn Patterson, Dr. Anna Woollams and Dr. Elisa Cooper for contributing to the real datasets utilised in this study. We are also very grateful for the valuable feedback from the anonymous reviewers that helped us improve the work.

## Appendix A The effect of leakage on multivariate connectivity

We can generalise the bivariate (two-ROI) example discussed in section 2.1.4 to multivariate methods for estimating the unique (partial) covariance between pairs of ROIs in a network of connections between three or more ROIs. In Fig. 1 d, consider a seed in the RMF (region Y), a target in the MTG (region Z) and a new region X within the leakage realm of MTG. Let us assume that the true source in Z co-varies with Y, but true connectivity between X and Y is zero. Let us further assume, for the sake of simplicity, that the whole network only consists of these three regions and Y does not receive/send leakage from/to any other ROIs. Therefore, considering the linear and time-unvarying effects of leakage, the estimated X and Z signals will be a linear combination of true signals at these regions (X’ and Z’ respectively) while the estimated Y activity equals the true source activity Y’ and can be written as:

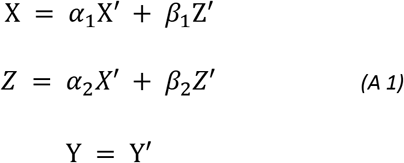

where α1 and β1 are the amount of leakage that X receives from itself and true Z’ source respectively and α_2_ and β_2_ are the amount of leakage that Z receives from true X’ source and itself respectively. Therefore, in the scenario outlined above, COV_X’Y’_=0 and in order for the partialising of covariance to overcome leakage, it should yield COV_XY|Z_=0.

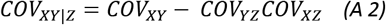

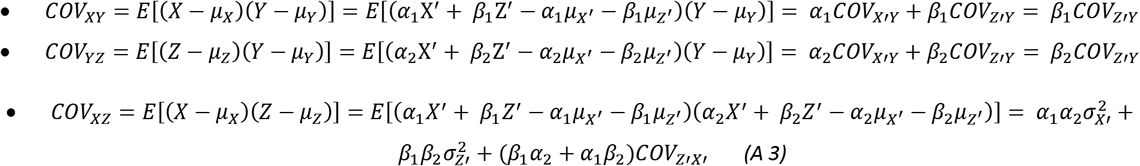

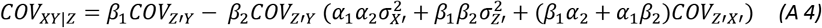

Therefore, COV_XY|Z_≠0. The only exceptional case is when the true source X’=μ_x’_ (i.e. inactive), β_1_=β_2_=1(i.e. Z and X are equally influenced by the leakage from Z), Z’ has unit variance and, thus. Even though the second condition (β_1_=β_2_=1) might be obviated using normalised measures of co-variation, the first and third conditions are unlikely to be true for the whole brain network analysis.

This argument, likewise for the bivariate methods, might be generalised to time-lagged connectivity measures (e.g. multivariate autoregressive modelling).

Even though the above examples argue that leakage cannot be resolved using non-zero-lag or multivariate connectivity measures, Equations A2–A4 show that quantifying leakage between ROIs (i.e. coefficients α_1_ α_2_ β_1_ β_2_) and combining them with multivariate connectivity measures might provide a more accurate reconstruction of whole brain networks using source reconstructed EEG/MEG data. In this study we concentrated on the former.

## Appendix B Simulations supplementary materials

### B.1. Brainnetome Atlas and coverage by modified parcellations

**Figure B. 1.**
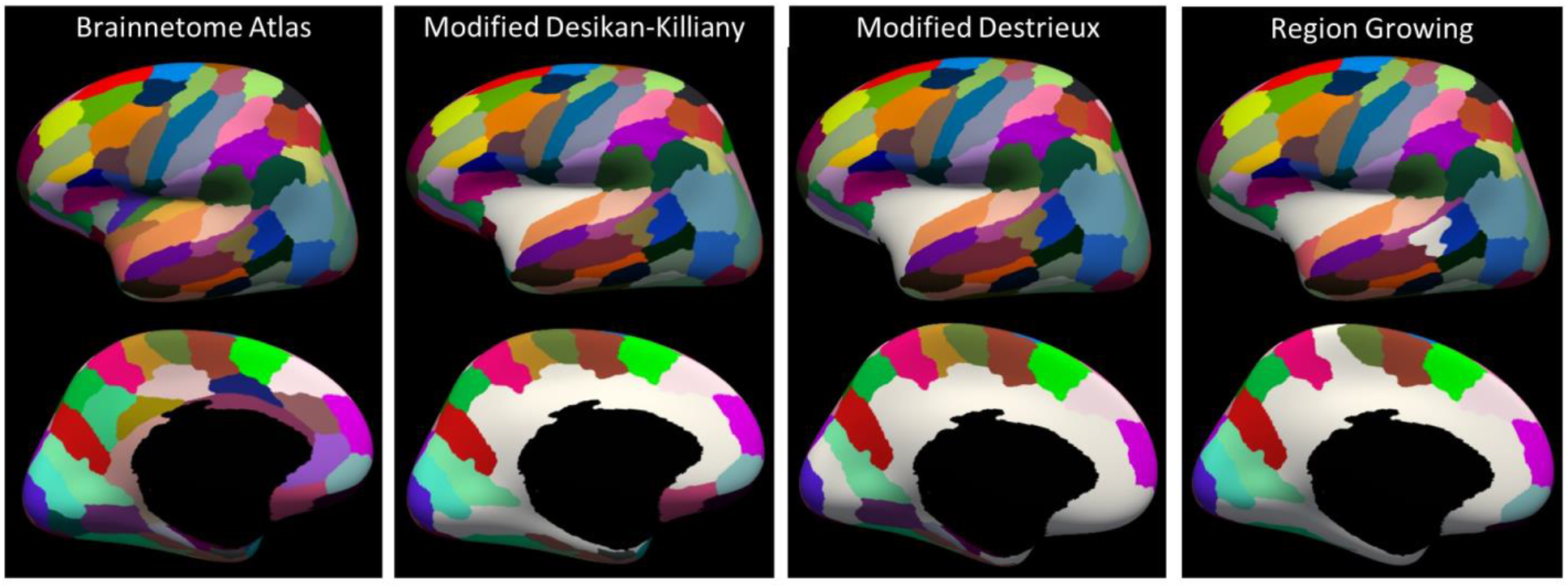
a) Brainnetome functional atlas from which the active nodes (ANs) in the brain are randomly drawn. parcels of the Brainnetome atlas that show any overlap with parcels of the modified a) Desikan-Killiany, b) Destrieux and c) Region Growing parcellations.

### B.2. Simulation results

In the results section, under question 2 section 4.4.2, we evaluated the performance of different parcellation algorithms using simulated networks with different number of ANs and connections. Here, we aim to observe the sensitivity of each parcellation to the areas covered by each parcellation. Motivations are brought in ‎4.4.2 and results are presented in Fig. B. 2 and Table B. 1

**Figure B 2.**
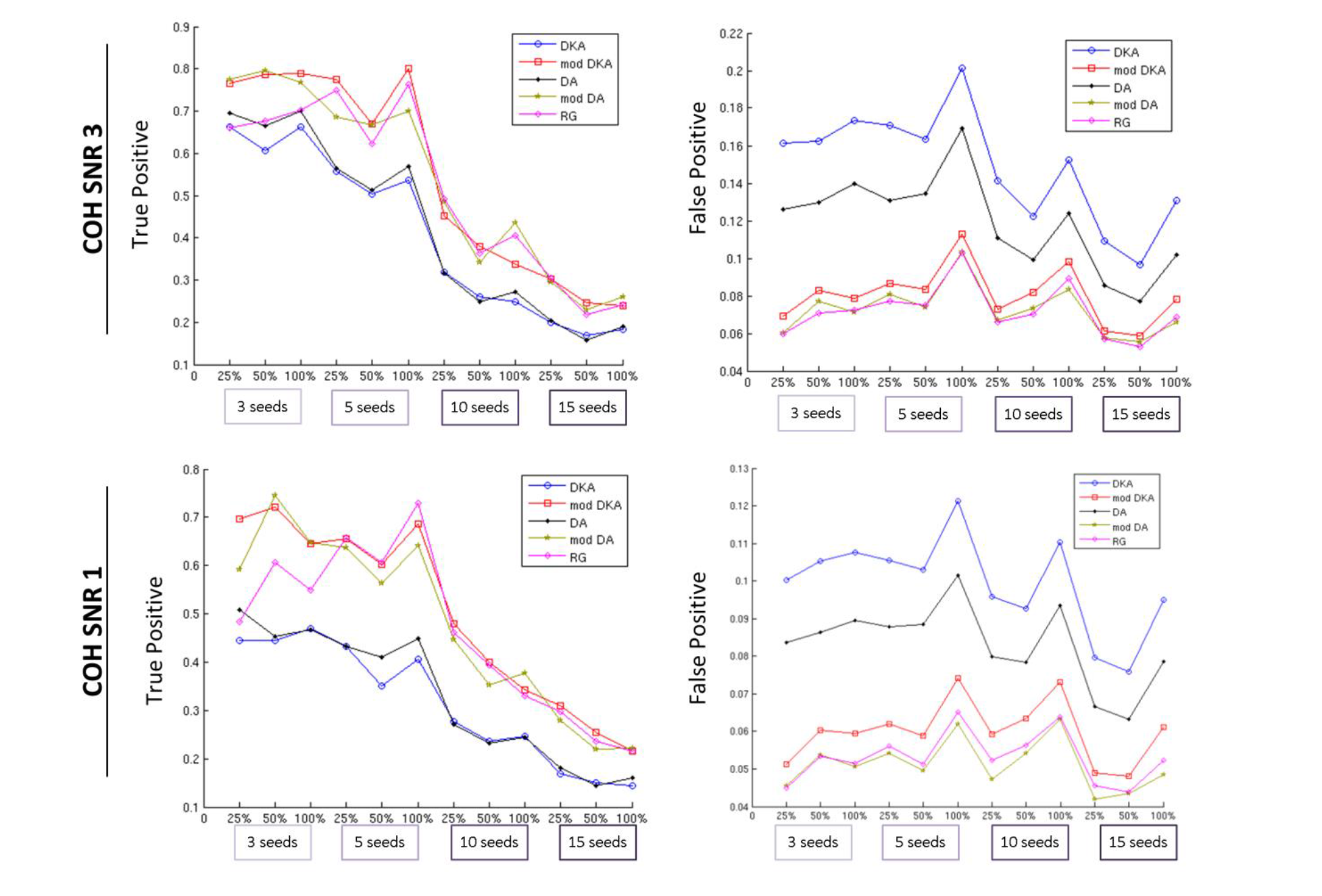
TPRs and FPRs before taking the missed connections due to no coverage of some parts of the cortex in adaptive parcellations into account. That is, we compared parcellation-specific ground truths (e.g. first row in Fig. 8) in the absence of leakage to the realistic networks in the presence of leakage. Left panel, TPRs: at SNR 3 (top), TPRs were reduced as the number of seeds/connections increased. For the anatomical parcellations, starting from ~0.7 true positives for 3 seeds, TPR was reduced to ~0.5 for 5 seeds, and then dropped sharply to 0.3 or less for 10 and 15. We found a similar trend for the modified parcellations, except that a) for all the seeds/connections, modified parcellations showed significantly higher TPRs than anatomical parcellations (c.f. Table B. 1 for details) and b) for 3 and 5 seeds, the TPR remained relatively constant at around 0.75 and then dropped to 0.4 or less for 10 seeds or more. We found no significant difference between the TPRs of the three modified parcellations. Both SNRs showed a similar trend, but (as elaborated in Table B. 1), the TPRs at SNR 1 where on average lower by approximately 10% and 6% compared to those at SNR 3, for anatomical and modified parcellations respectively. Right panel, FPRs: FPRs are the same as Fig. 9 and are presented here for the sake of completeness.

**Table B 1.**
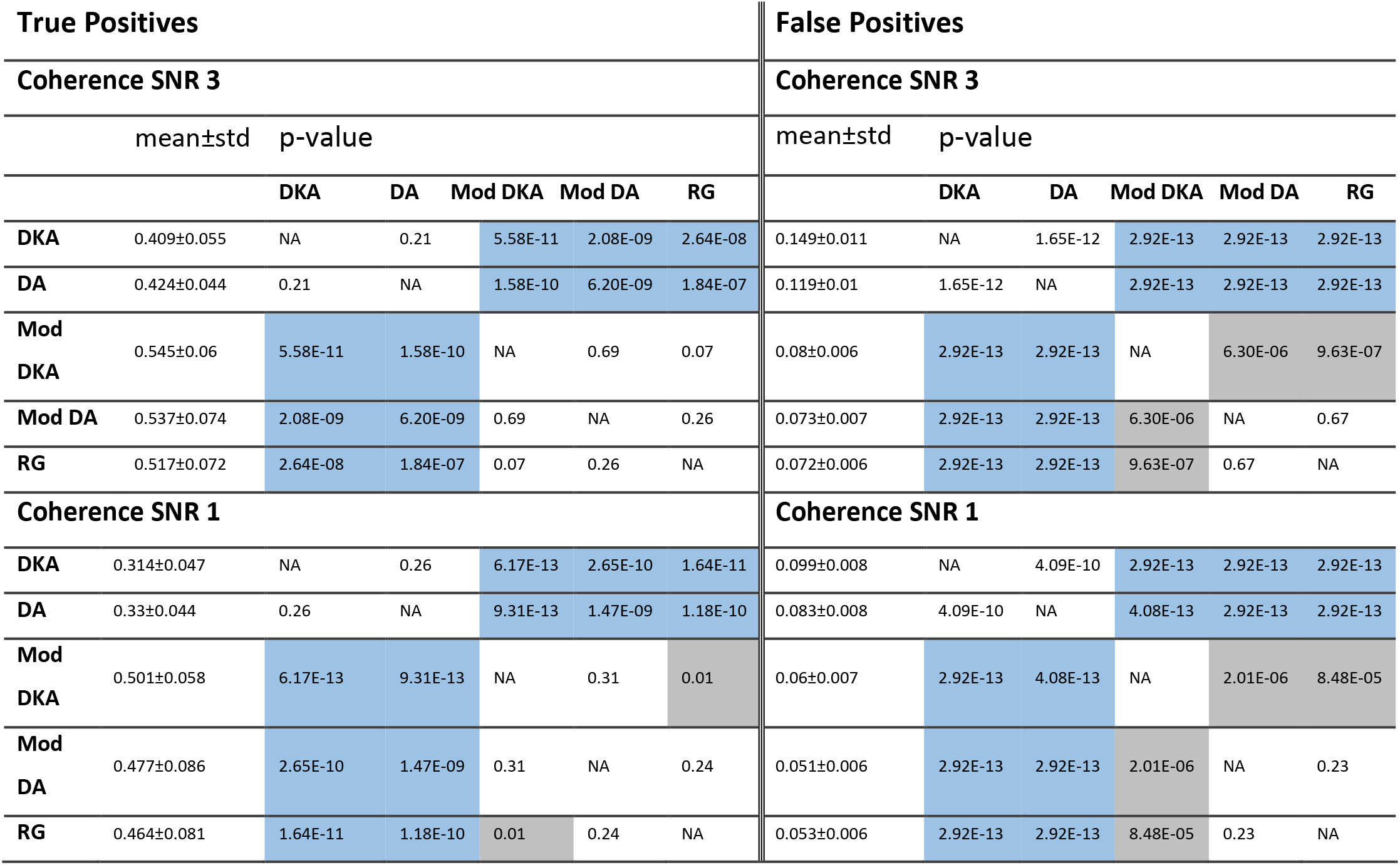
Statistical comparison of average TPRs and FPRs of anatomical and modified parcellation without considering the uncovered ANs in the modified parcellations. Blue shading: adaptive significantly improved compared to anatomical. Grey shading: Significant differences among the adaptive parcellations.

## Appendix C Initial results of the parcellation algorithms

**Fig. C. 1.**
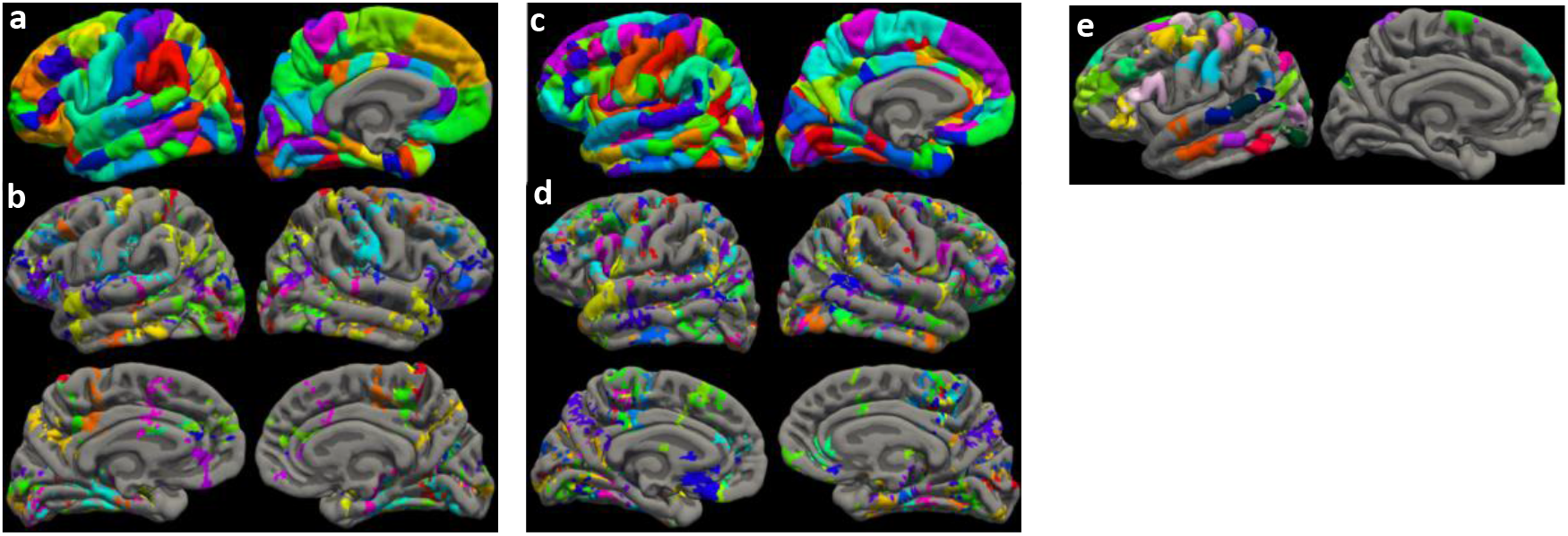
shows the intitial split and merged parcels which were input to the final parcellation procedure. Figure C 1 Initial results of the parcellation algorithms. a) split and b) merged parcels from the Desikan-Killiany Atlas. The primary splitting procedure for this parcellation resulted in 194 split parcels and merging procedure resulted in 122 merged parcels; these sets of parcels formed an intermediate parcellation that was input into the final homogeneity check for final vertex assignment (section 3.3.2). From this figure, it can be seen that larger parcels (e.g. pre-/post-central and temporal regions) were split into several sub-parcels. Additionally, vertices that are located at the intersection of adjacent parcels were typically clustered together to form merged parcels. While some of these clusters survive as new parcels in the final parcellation (Fig 4a), others are removed, leaving gaps between the neighbouring parcels which results in a sparse sampling of the cortex to maximise the distinguishability in the final parcellation. c) split and d) merged parcels from Destrieux Atlas. The initial splitting procedure for this parcellation resulted in 428 parcels and merging procedure in 280 extra parcels (overall 708 parcels) compared to 316 parcels for the Desikan-Killiany atlas. e) “Created” parcels from the region growing algorithm. These created parcels were mirrored to the right hemisphere using MNI coordinates.

